# Tn*Smu1* is a functional integrative and conjugative element in *Streptococcus mutans* that when expressed causes growth arrest of host bacteria

**DOI:** 10.1101/2022.08.23.505017

**Authors:** Lisa K. McLellan, Mary E. Anderson, Alan D. Grossman

**Affiliations:** Department of Biology Massachusetts Institute of Technology Cambridge, Massachusetts 02139

**Keywords:** horizontal gene transfer, conjugation, mobile genetic element, integrative and conjugative element, *Streptococcus mutans*

## Abstract

Integrative and conjugative elements (ICEs) are major drivers of horizontal gene transfer in bacteria. They mediate their own transfer from host cells (donors) to recipients and allow bacteria to acquire new phenotypes, including pathogenic and metabolic capabilities and drug resistances. *Streptococcus mutans,* a major causative agent of dental caries, contains a putative ICE, Tn*Smu1*, integrated at the 3’ end of a leucyl tRNA gene. We found that Tn*Smu1* is a functional ICE, containing all the genes necessary for ICE function. It excised from the chromosome and excision was stimulated by DNA damage. We identified the DNA junctions generated by excision of Tn*Smu1*, defined the ends of the element, and detected the extrachromosomal circle. We found that Tn*Smu1* can transfer from *S. mutans* donors to recipients when co-cultured on solid medium. The presence of Tn*Smu1* in recipients inhibited successful acquisition of another copy and this inhibition was mediated, at least in part, by the likely transcriptional repressor encoded by the element. Using microscopy to track individual cells, we found that activation Tn*Smu1* caused an arrest of cell growth. Our results demonstrate that Tn*Smu1* is a functional ICE that affects the biology of its host cells.

## Introduction

Horizontal gene transfer (HGT) is a driving force in microbial evolution, allowing bacteria to acquire new traits and phenotypes from other bacterial lineages. Biofilms, including dental plaque, are hot spots for HGT, and HGT is well documented in the oral microbiome (Jones et al., 2021; Lunde et al., 2021; Olsen et al., 2013; Roberts et al., 1999, 2001). Further, oral bacteria can cause major health issues. For example, *Streptococcus mutans,* a major causative agent of dental caries, acts as a reservoir for antibiotic resistance genes and mobile genetic elements within the oral microbiome and can be a causative agent of infective endocarditis (Lunde et al., 2021; Nomura et al.; Olsen et al., 2013).

Much is known about quorum sensing and HGT through natural competence in *S. mutans* (reviewed in (Shanker and Federle, 2017)). Other types of HGT are mediated by mobile genetic elements that are often found integrated in the genome of the host organism. These can have a broad host range and some can mediate genetic exchange between distantly related organisms that may be unable to exchange DNA through transformation. Conjugative elements and bacteriophage are mobile genetic elements that can mediate their own transfer from a host (donor) bacterium to a recipient.

Integrative and conjugative elements (ICEs) represent the most prevalent type of conjugative element and are found in all major clades of bacteria, including oral and cavity-causing bacteria (Guglielmini et al., 2011; Lunde et al., 2021; Roberts and Mullany, 2009, 2011; Roberts et al., 2001). Integrative and conjugative elements mediate their own transfer from a host cell to a recipient and often contain cargo genes that confer beneficial phenotypes to the host cells. Phenotypes conferred by ICEs include: virulence, symbiosis, metabolic functions, and drug resistances (reviewed in (Johnson and Grossman, 2015)). ICEs are major contributors to the spread of antibiotic resistance determinants among multidrug-resistant pathogens and are notable in the human microbiome and other microbial communities. However, fundamental aspects of ICE biology within the complex communities of dental plaque remain unknown.

Although their size, regulation, and interactions with host cells can vary considerably, ICEs generally follow a standard lifecycle (reviewed in (Auchtung et al., 2016; Botelho and Schulenburg, 2021; Wozniak and Waldor, 2010), among others). Briefly, ICEs typically reside integrated in the chromosome of a bacterial host. Under certain conditions or perhaps stochastically, ICE gene expression becomes active and the ICE excises from the host chromosome to form an extrachromosomal circular dsDNA molecule. This is then processed by ICE and host-encoded proteins to generate a nucleoprotein complex containing linear single-stranded ICE DNA. Element DNA (as linear ssDNA) can then be transferred to a new host through the element-encoded type IV secretion system (T4SS). Once in the recipient, the linear ssDNA circularizes and is converted into dsDNA. The circular dsDNA can then be integrated into the recipient genome through the action of an ICE-encoded recombinase, generating a stable transconjugant (recipient that has stably acquired a copy of an ICE).

While integrated in the host chromosome, expression of most of the genes required for the ICE lifecycle are repressed. In many ICEs, this is controlled by a transcriptional repressor. Upon activating signals, the repressor is inactivated, sometimes via proteolytic cleavage by an element-encoded anti-repressor (Bose et al., 2008). This allows the genes required for ICE transfer to be expressed and the ICE lifecycle to continue.

Throughout its lifecycle, the interactions between an ICE and its host cell are complex. ICEs often benefit their host cells through associated cargo genes that confer specific phenotypes (mentioned above). However, ICEs can also manipulate host development, growth, and viability (Beaber et al., 2004; Bean et al., 2022a; Jones et al., 2021; Pembroke and Stevens, 1984; Reinhard et al., 2013).

Tn*Smu1* is a putative ICE found in some isolates of the prototypic oral pathogen *S. mutans.* (Ajdic et al., 2002; Bi et al., 2012). Tn*Smu1* was recognized bioinformatically based on comparisons to the conjugation machinery and DNA processing machinery of two well studied ICEs: Tn*916,* the first described ICE discovered through its ability to spread tetracycline resistance through clinical isolates of *Enterococcus* (Franke and Clewell, 1981a, 1981b), and ICE*Bs1,* a well-studied ICE in *Bacillus subtilis* (Auchtung et al., 2007, 2016; Jones et al., 2021; Lee and Grossman, 2007; Lee et al., 2007).

We found that Tn*Smu1* is a functional ICE that is capable of undergoing a complete ICE life cycle. We demonstrate that Tn*smu1* can excise from its host chromosome to form a circular dsDNA molecule, transfer to recipient cells, and integrate into the chromosome to generate stable transconjugants. Using PCR-based assays, we identified the integration site, defined the ends of the element, and detected the extrachromosomal circle. We found that DNA damage modulates the excision of T*nSmu1.* Further, by co-culturing donors and recipients on a solid medium, we found that Tn*Smu1* can transfer from *S. mutans* donors to recipients. However, the presence of Tn*Smu1* in recipients prevented successful acquisition of another copy of the element, at least in part, through a repressor-mediated immunity mechanism analogous to that of ICE*Bs1* (Auchtung et al., 2007; Bose et al., 2008). Using a fluorescent reporter in Tn*Smu1*, we were able to visualize individual cells in which Tn*Smu1* had become active. Most of the cells with an active element stopped growing. Together, our results illuminate the functionality of Tn*Smu1* and demonstrate that Tn*Smu1* affects the physiology of its host cells by halting host cell growth. As our work was nearing completion, an independent study identified the putative repressor of Tn*Smu1* and reported that the element can be activated to excise from the chromosome by conditional inactivation of the repressor and also by DNA damage (King et al., 2022).

## Results

### Comparison of Tn*Smu1* to Tn*916* and ICE*Bs1*

Tn*Smu1* (Fig. 1) is a putative ICE found in the prototypic oral pathogen *S. mutans* UA159 (Ajdic et al., 2002; Bi et al., 2012). Tn*Smu1* was recognized bioinformatically based on comparisons to genes from the ICE Tn*916* that are needed for conjugation and DNA processing (Table 1) (Ajdic et al., 2002; Roberts and Mullany, 2009, 2011). *S. mutans* appears to be the primary host of Tn*Smu1* (Fig. S1), although remnants or fragments of the element are found in other Streptococcal species.

**Figure 1.**
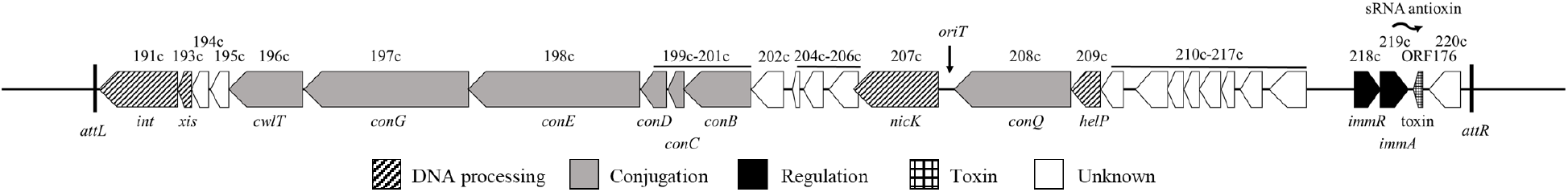
Genetic map of Tn*Smu1.* Open reading frames are indicated by horizontal arrows. Gene names are abbreviated to include only the number designation (i.e., 191c indicates *smu_191c*). The name of the homologous ICE*Bs1* gene is written below. Thick vertical black lines indicate attachment sites *attL* and *attR* at the ends of Tn*Smu1*. The putative origin of transfer (*oriT*) is indicated with a vertical arrow. Putative gene function is indicated by pattern and color: genes of unknown color (white), genes encoding the type 4 secretion system (grey), DNA processing (diagonal stipes), and regulation (black). There is a toxin (grid pattern) and small RNA antitoxin (wavy horizontal arrow) encoded in the region between *smu_217c* and *smu_218c* (Koyanagi and Lévesque, 2013).

**Table 1.** Similarity between ICEs Tn*Smu1,* ICE*Bs1,* and Tn*916*.

| Tn*SmuI*<br>gene<sup>1</sup> | Proposed<br>name<sup>2</sup> | % similarity (% identity)<sup>3</sup> |  |  |  | Vir homolog/function/comments<sup>4</sup> |
| --- | --- | --- | --- | --- | --- | --- |
|  |  | ICE*BsI* |  | Tn*916* |  |  |
| *smu_191c* | *int* | *int* | 35.8 (19.0) | *int* | 37.6 (21.5) | tyrosine recombinase |
| *smu_193c* | *xis* | *xis* | 31.6 (14.5) | *xis* | 34.2 (15.1) | excisionase/recombination<br>directionality factor |
| *smu_196c* | *cwlT* | *cwlT* | 25.6 (17.5) | *orf14* | 32.7 (19.3) | VirB1-like, cell wall hydrolase |
| *smu_197c* | *conG* | *conG* | 26.0 (15.1) | *orf15* | 29.2 (15.8) | VirB6-like, transmembrane protein |
| *smu_198c* | *conE* | *conE* | 46.2 (28.1) | *orf16* | 38.5 (22.7) | VirB4-like, AAA+ ATPase |
| *smu_199c* | *conD* | *conD* | 33.2 (16.8) | *orf17* | 24.7 (15.9) | VirB3-like, transmembrane protein |
| *smu_200c* | *conC* | *conC* | 26.0 (12.5) | *orf19* | 29.2 (19.8) | transmembrane protein |
| *smu_201c* | *conB* | *conB* | 37.0 (21.4) | *orf13* | 36.8 (18.4) | VirB8-like, transmembrane protein |
| *smu_207c* | *nicK* | *nicK* | 40.4 (24.2) | *orf20* | 46.5 (25.6) | relaxase |
| *smu_208c* | *conQ* | *conQ* | 44.2 (26.7) | *orf21* | 42.6 (26.7) | VirD4-like, AAA+ ATPase; coupling<br>protein |
| *smu_209c* | *helP* | *helP* | 40.8 (24.3) | *orf22*<br>*orf23* | 38.3 (22.7)<br>30.8 (22.6) | helicase processivity factor |
| *smu_218c* | *immR* | *immR* | 51.9 (31.3) | n/a |  | repressor; DNA binding protein |
| *smu_219c* | *immA* | *immA* | 30.2 (16.1) | n/a |  | anti-repressor; metalloprotease, cleaves<br>ImmR |

<sup>1</sup>Tn*Smu1* gene names from (Ajdic et al., 2002)
<sup>2</sup>Proposed name based on similarity to known genes. Uses ICE*Bs1* gene names where appropriate based on conservation (Auchtung et al., 2016).
<sup>3</sup>Calculated by EMBOSS Needle pairwise global sequence alignment (Needleman and Wunsch, 1970)
<sup>4</sup>Function based on similarity to characterized proteins (Auchtung et al., 2016; Fronzes et al., 2009)

In addition to similarity to genes in Tn*916*, many genes in Tn*Smu1* are similar to those in ICE*Bs1* from *B. subtilis* (Table 1) (Auchtung et al., 2016). We analyzed sequence similarities between genes in Tn*Smu1*, ICE*Bs1*, and Tn*916* and found that Tn*Smu1* is predicted to contain most, if not all components that are needed for the ICE lifecycle (Fig. 1, Table 1).

#### Type IV secretion system

We identified all the known components of the T4SS typical of Gram-positive bacteria. The membrane channel of the T4SS is likely composed of ConB, ConC, ConD, and ConG (ConB_ICE*Bs1*_, ConC _ICE*Bs1*_, ConD _ICE*Bs1*_, and ConG _ICE*Bs1*_ and ORF13_Tn*916*_, ORF19_Tn*916*_, ORF17_Tn*916*_, and ORF15_Tn*916*_, for ICE*Bs1* and Tn*916* respectively) (Auchtung et al., 2016; Ciric et al., 2013; Leonetti et al., 2015; Roberts and Mullany, 2009). All of these proteins are predicted to encoded a specific number of transmembrane domains and the approximate size of the proteins is conserved throughout T4SSs (Auchtung et al., 2016; Berkmen et al., 2010; Leonetti et al., 2015). Based on homology, size, and predicted transmembrane localization, we precited *smu_201c*, *smu_200c*, *smu_199c*, and *smu_197c* to encode ConB_Tn*Smu1*_, ConC _Tn*Smu1*_, ConD _Tn*Smu1*_, and ConG_Tn*Smu1*_ respectively (Table 1).

ConE (ConE_ICE*Bs1*_ and ORF16_Tn*916*_) is one of two conserved ATPases found in T4SSs of Gram-positive bacteria (Auchtung et al., 2016; Fronzes et al., 2009) that is required for conjugation (Auchtung et al., 2007; Berkmen et al., 2010; Iyer et al., 2004; Leonetti et al., 2015). *smu_198c* encodes a gene product that is 46.2% similar to ConE_ICE*Bs1*_ and 38.5% similar to ORF16_Tn*916*_ (Table 1), and thus we infer SMU_198c is ConE_Tn*Smu1*_.

The coupling protein, ConQ_ICE*Bs1*_ and ORF21_Tn*916*_, binds the DNA protein complex (relaxasome) and ‘delivers’ or ‘couples’ it to the T4SS for transfer into a recipient cell. Key features of coupling proteins from T4SSs of Gram-positive bacteria include two transmembrane helices in the N-terminal domain and a C-terminal cytoplasmic ATPase domain (Alvarez-Martinez and Christie, 2009; Auchtung et al., 2016). ConQ_Tn*Smu1*_ (SMU_208c) shares 44.2% and 42.6% similarity with ConQ_ICE*Bs1*_ and ORF21_Tn*916*_ respectively and is predicted to have the same structural features found in other coupling proteins (Table 1).

#### Cell wall hydrolase

Cell wall hydrolases are critical components of conjugative elements from Gram-positive bacteria (Auchtung et al., 2016; Bhatty et al., 2013). SMU_196c contains a phage tail lysozyme domain (E-value 2.20e-30 BLASTp), used by phage to degrade the cell wall peptidoglycan layer (Xiang et al., 2008). It also includes an amidase domain (E-value 1.23e-48 BLASTp). Structural predictions through Phyre2 matched the structure of SMU_196c to a cell wall hydrolase produced by *Staphylococcus aureus* (N-acetylmuramoyl-L-alanine amidase 2 from *S. aureus,* 100% confidence, 76% coverage). Therefore, we determined this is likely the cell wall hydrolase for Tn*Smu1* and we infer SMU_196c is CwlT_Tn*Smu1*_.

#### DNA relaxase, origin of transfer (*oriT*), and helicase processivity factor

After excision from the chromosome, the double stranded circular DNA is nicked and unwound for a single strand of DNA to be transferred through the conjugation machinery. Nicking occurs at the origin of transfer (*oriT*) through the action of a DNA relaxase (or nickase) (NicK_ICE*Bs1*_ and ORF20_Tn*916*_) (Auchtung et al., 2016; Lee and Grossman, 2007; Rocco and Churchward, 2006). At least several ICEs undergo autonomous rolling-circle replication (Johnson and Grossman, 2015; Lee et al., 2010; Wright and Grossman, 2016). Like conjugation, replication initiates from *oriT* by the relaxase and also requires unwinding of the duplex DNA. In *B. subtilis*, DNA unwinding is mediated by the host-encoded DNA translocase PcrA (Lee et al., 2010; Petit et al., 1998; Thomas et al., 2013). For ICE*Bs1* and Tn*916*, a helicase processivity factor (HelP_ICE*Bs1*_ and *orf22* and *orf23* for Tn*916*) is required to facilitate DNA unwinding (Thomas et al., 2013). The genes encoding the processivity factors are immediately upstream of and in a cluster with *conQ* (coupling protein), *oriT*, and *nicK* (relaxase), in that order (Thomas et al., 2013) (Fig. 1).

**i) *nicK*_Tn_*_Smu1_*.** *smu_207c* in Tn*Smu1* encodes a product that is 40.4% and 46.5% similar to NicK_ICE*Bs1*_ and ORF20_Tn*916*_, respectively (Table 1). Similar to the location in ICE*Bs1* and Tn*916*, it is immediately downstream from the gene encoding the coupling protein (Fig. 1). Based on the sequence similarity and location, we infer that SMU_207c is NicK_Tn*Smu1*_.

**ii) *oriT*_Tn_*_Smu1_*.** We identified a sequence in Tn*Smu1* that is identical in 20 of 24 bp to *oriT* from ICE*Bs1* and Tn*916* (Fig. 2A) (Lee and Grossman, 2007)*. oriT*_Tn*Smu1*_ contains an inverted repeat that is characteristic of many *oriT*s and is located just upstream of *nicK* (Fig. 1). Based on the sequence and location, we infer that this is *oriT*_Tn*Smu1*_.

**Figure 2.**
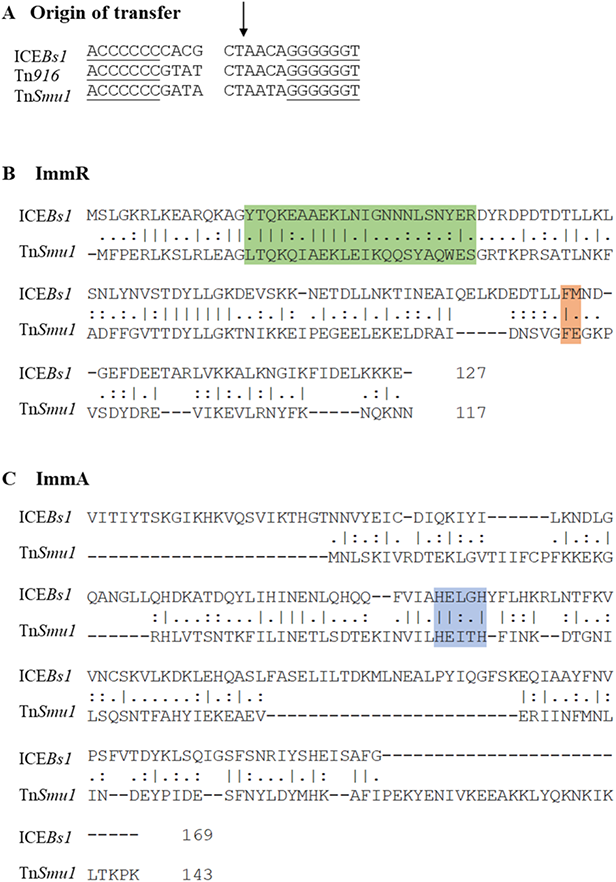
Alignments of *oriT*, ImmR, and ImmA from Tn*Smu1,* ICE*Bs1,* and Tn*916*. **A.** Alignment of the putative Tn*Smu1* origin of transfer (*oriT*) to ICE*Bs1* and Tn*916.* The *nic* site of ICE*Bs1* and Tn*916* is indicated by a vertical arrow. Inverted repeats are indicated by lines under the sequence. **B, C.** Global alignments of ImmR (**B**) and ImmA (**C**) from ICE*Bs1* and Tn*Smu1* as calculated by EMBOSS Needle pairwise alignment (Needleman and Wunsch, 1970). “|” represents a matching amino acid; “:” represents amino acids with strongly similar properties; “.” represents amino acids with weakly similar properties; “-” represents a gap (Rice et al., 2000). **B**. For ImmR, amino acids boxed in green represent a conserved helix-turn-helix motif (Bose et al., 2008; Oppenheim et al., 2005) (predicted by GYM 2.0 (Gao et al., 1999; Narasimhan et al., 2002)). Amino acids boxed in orange represent the cleavage site where ImmR_ICE*Bs1*_ is cleaved by ImmA_ICE*Bs1*_ (Bose et al., 2008). **C.** For ImmA, amino acids boxed in blue indicate the characteristic HEXXH motif found in many zinc-dependent metalloproteases (Fujimura-Kamada et al., 1997).

**iii) *helP*_Tn_*_Smu1_*.** *smu_209c* in Tn*Smu1* encodes a product that is 40.8% similar to the helicase processivity factor HelP_ICE*Bs1*_ and 38.3% and 30.8% similar to Orf22 and Orf23, the two HelP homologs from Tn*916* (Table 1). Additionally, it is located immediately upstream from the gene (*conQ*) that encodes the coupling protein (Fig. 1). Based on the location and similarities, we infer that SMU_209c is HelP_Tn*Smu1*_.

#### Integrase and excisionase

Integrating elements typically utilize a recombinase, often called an integrase (Int), that is required for integration into and excision from a host chromosome. Integrating elements also utilize a recombination directionality factor, also called an excisonase (Xis), in combination with Int, for excision from the host chromosome (Grindley et al., 2006; Hirano et al., 2011).

*smu_191c* encodes a product that is 35.8% and 37.6% similar to the tyrosine recombinases (Int) encoded by ICE*Bs1* and Tn*916*, respectively (Table 1). Integrases (recombinases) are often encoded at or near an end of an ICE (Cury et al., 2017). Based on the location and protein similarities, we infer that SMU_191c is Int_Tn*Smu1*_.

*smu_193c* encodes a gene product that is 31.6% similar and 34.2% similar to Xis_ICE*Bs1*_ and Xis_Tn*916*_ (Table 1). Excisionases (Xis) are typically small, highly charged proteins often with little sequence similarity to other excisionases (Auchtung et al., 2016; Lee et al., 2007). *smu_193c* encodes is a small, basic protein and thus we infer that SMU_193c is likely Xis_Tn*Smu1*_.

#### Regulatory proteins (ImmA/ImmR)

Tn*Smu1* has two genes, *smu_218c* and *smu_219c* that encode products similar to the repressor (ImmR) and anti-repressor and metalloprotease (ImmA) encoded by ICE*Bs1*. SMU_218c and SMU_219c are 51.9% and 30.2% similar to ImmR_ICE*Bs1*_ and ImmA_ICE*Bs1*_, respectively (Fig. 2B, C) and we infer that the Tn*Smu1* products are ImmR_Tn*Smu1*_ (SMU_218c) and ImmA_Tn*Smu1*_ (SMU_219c). For ICE*Bs1*, when activated, the protease ImmA cleaves ImmR to inactivate it, thereby causing de-repression of the element (Auchtung et al., 2007; Bose et al., 2008). Homologs of ImmR and ImmA are also encoded by many phages (Bose et al., 2008; Lucchini et al., 1999). ImmR_Tn*Smu1*_ contains a conserved a helix-turn-helix motif, a typical DNA binding domain for phage-like repressors (Bose et al., 2008). It also has a conserved phenylalanine that may be the putative cleavage site (Fig. 2B) (Bose et al., 2008). That ImmR_Tn*Smu1*_ acts as the repressor of Tn*Smu1* is further supported by finding that it appears essential in Tn-seq studies (Shields et al., 2018), as would be expected if deletion of the gene caused de-repression and excision of the element, thereby deleting the entire element from the chromosome. Further, ImmA_Tn*Smu1*_ contains a characteristic HEXXH motif found in many zinc-dependent metalloproteases (Fig. 2C) (Fujimura-Kamada et al., 1997).

Based on the presence of the regulatory genes and the essential components of an ICE, we expected that Tn*Smu1* is a functional ICE. Below we describe experiments demonstrating that Tn*Smu1* is indeed a functional element: it can excise from the host chromosome, transfer to a new host, and integrate into the chromosome of the new host to generate a stable transconjugant.

### Excision of Tn*Smu1* from the host chromosome

When Tn*Smu1* is integrated in the host cell chromosome, there are left and right attachment sites, *attL* and *attR*, respectively, that demarcate the junctions between Tn*Smu1* and the host chromosome (Fig. 3A). If capable of and upon excision, there would be a single attachment site in the element, *att*Tn*Smu1*, where the element recombined to form an extrachromosomal circle. There would also be an attachment site in the bacterial chromosome, *attB*, that represents the fusion of chromosomal sequences that had been interrupted by insertion of the element (Fig. 3A).

**Figure 3.**
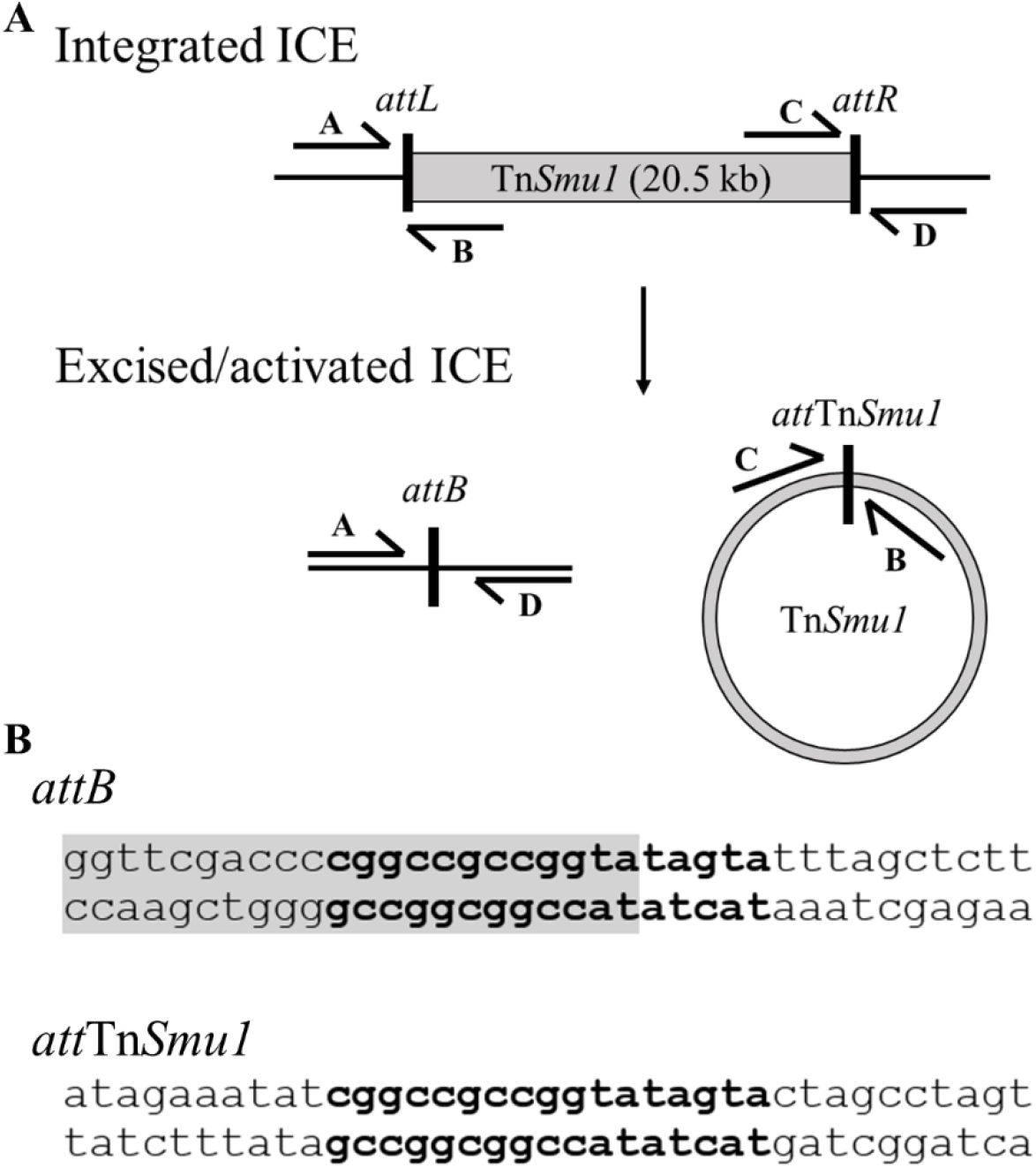
Cartoon of Tn*Smu1* inserted in the chromosome and products and sequences after excision. **A.** Cartoon of Tn*Smu1* inserted in the chromosome and the products after excision. A set of four primers (labeled A, B, C, D) are able to detect the junctions between the host chromosome and left (*attL*; primers A+B) and right (*attR*; primers C+D) ends of Tn*Smu1*; the host attachment site without insertion of Tn*Smu1* (*attB*; primers A+D) and the excised Tn*Smu1* circle (*att*Tn*Smu1*; primers B+C). **B.** Genome sequence of *attB* and *att*Tn*Smu1*. The 17 bp recombination site is in bold. The 3’ end of the leucyl tRNA (*smu_t33*) that overlaps the recombination site is highlighted in gray.

*attL* had been predicted based on the increase in A+T content throughout the element that is characteristic of horizontally acquired DNA, and the presence of nearby genes encoding putative recombinases (*smu_191c)* (Ajdic et al., 2002; Bi et al., 2012). We noticed that Tn*Smu1 attL* was in *smu_t33*, encoding a leucyl tRNA. ICEs are often found integrated into tRNA genes and typically do not disrupt the gene (Burrus and Waldor, 2004; Burrus et al., 2002). For example, ICE*Bs1* and ICE*Hin1056* from *B. subtilis* and *Haemophilus influenzae*, respectively, are both found integrated in a tRNA gene (Dimopoulou et al., 2002; Lee et al., 2007). The location of *attR* of Tn*Smu1* had been predicted to be either after *smu_226c* or *smu_209c,* based on A+T content, the presence of genes encoding an integrase, relaxase, and-or type IV secretion system, and-or the presence of a large intergenic region (Ajdic et al., 2002; Bi et al., 2012).

To test Tn*Smu1* excision, we designed primer sets upstream and downstream of the predicted ends (*attL* and *attR*) that would only produce a PCR product upon excision (Fig. 3A). We also designed primers internal to Tn*Smu1* but oriented outwards to detect the circularized element (Fig. 3A). We purified DNA from stationary phase cultures of *S. mutans* and were unable to detect excision of Tn*Smu1* using these initial primer sets. We found that this inability to detect excision of Tn*Smu1* was because *attR* was different from what was postulated initially and not because the element failed to excise.

We tested other primer pairs, keeping the primer near *attL* constant and changing the location of the primers near the putative *attR*. We identified PCR primers that produced products that, when sequenced, allowed us to define *attL, attR, attB*, and *att*Tn*Smu1* (Fig. 3A). Based on these sites, we conclude that Tn*Smu1* is an approximately 20 kb element that extends from open reading frames *smu_191c* through *smu_220c* (Fig. 1). The sequences of the various sites also allowed us to identify a 17 bp sequence present in both *attB* and *att*Tn*Smu1* that is likely the recombination site used by the tyrosine recombinase encoded by *int* (*smu_191c*) in Tn*Smu1* (Fig. 3B). This is consent with excision noted by King, *et al*. when examining read coverage in an *immR*_Tn*Smu1*_ mutant (King et al., 2022). Together, these data demonstrate that Tn*Smu1* is capable of excision from the chromosome of host cells.

### Tn*Smu1* excision increases in response to DNA damage and on solid media

Because Tn*Smu1* has homologs of *immA* and *immR* from ICE*Bs1* (Fig. 2B/C, Table 1) and ICE*Bs1* is de-repressed by DNA damage, we postulated that DNA damage might also de-repress Tn*Smu1*. Therefore, we measured excision of Tn*Smu1* after addition of mitomycin C (MMC) to cells to induce DNA damage. We purified DNA from cells treated and untreated with MMC and performed qPCR to detect the presence of *attB* (generated upon excision of Tn*Smu1*). We normalized excision to a nearby chromosomal locus (*ilvB*). We found that Tn*Smu1* had excised in ∼0.1 - 0.2% of cells (∼1-2×10^-3^ *attB/ ilvB)* after treatment with MMC for 2-4 hours. This is ∼20-30-fold greater than that in the absence of MMC (∼0.006%, ∼6×10^-5^ *attB/ ilvB*) (Fig. 4A). These results indicate that excision of Tn*Smu1* is induced following DNA damage.

**Figure 4.**
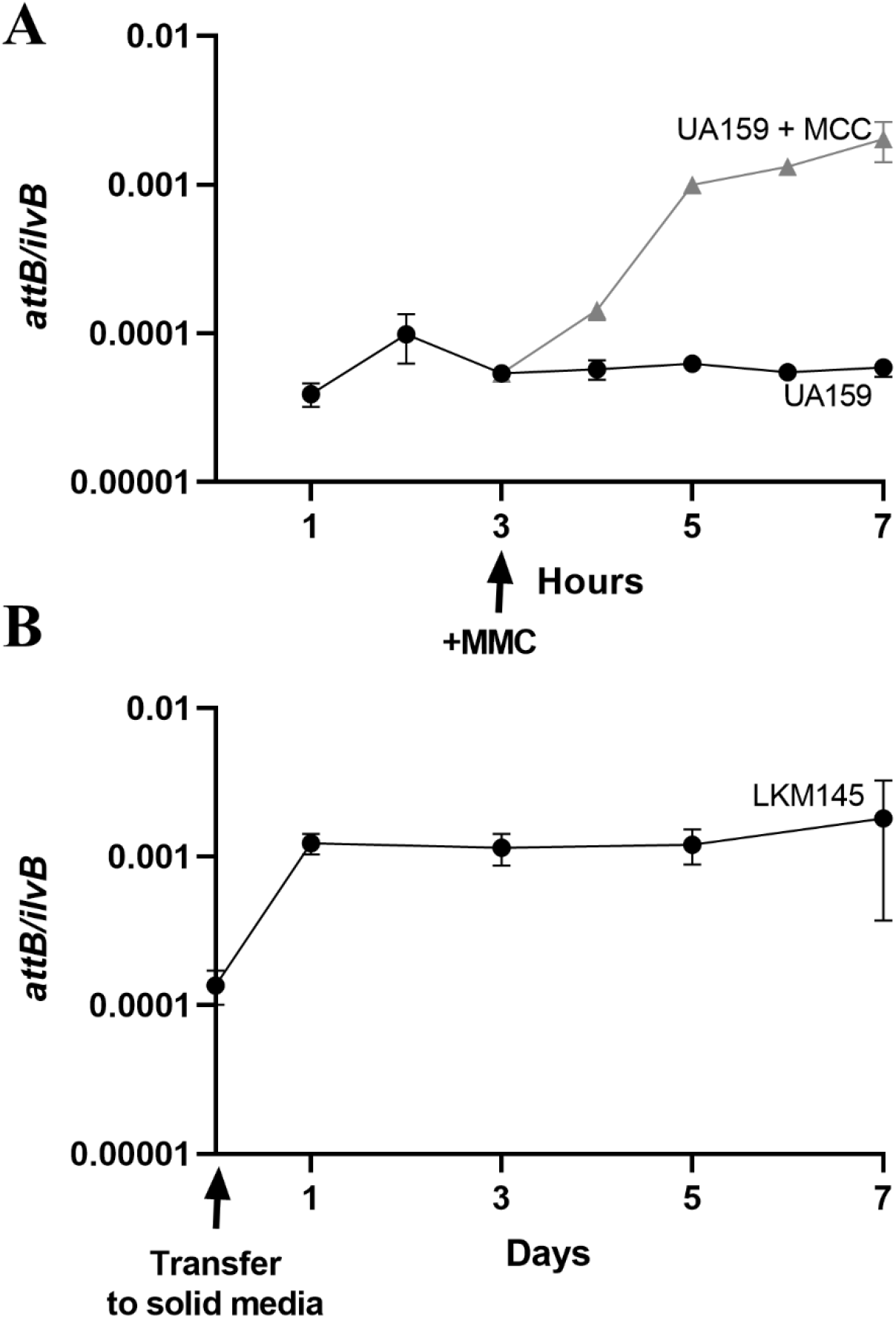
Tn*Smu1* excision increases in response to DNA damage and on solid medium. Excision of Tn*Smu1* was measured by qPCR to detect *attB* (primers corresponding to A+D indicated in Fig. 3). The proportion of cells containing excised Tn*Smu1* was calculated by normalizing *attB* to a nearby chromosomal locus (*ilvB*). Data presented are averages from three or more independent experiments with error bars depicting standard error of the mean. Error bars could not always be depicted due to the size of each data point. **A.** *S. mutans* UA159 cells were grown in liquid medium for seven hours. After 3 hours of growth, cells were either left untreated (black circles) or treated with 1 µg/ml mitomycin C (MMC; gray triangles; time of addition indicated by black arrow below the x-axis). Samples were harvested at the indicated times to measure excision. **B.** *S. mutans* strain LKM145 (Tn*Smu1-tet*) was grown in liquid medium to mid-exponential phase, pelleted, and resuspended at a low density. Cells were then spotted and grown on solid medium for 1, 3, 5, or 7 days in anaerobic conditions and samples were taken at the indicated times (days) to measure excision.

It also appeared that Tn*Smu1* underwent autonomous replication following excision. Both ICE*Bs1* and Tn*916* undergo autonomous rolling-circle replication following excision (Lee et al., 2010; Wright and Grossman, 2016). This is most easily detected by measuring the relative copy number of the circular element (*att*Tn*Smu1*) to that of the empty chromosomal site (*attB*). We found that during exponential growth, there were approximately 2-5 extrachromosomal copies of Tn*Smu1* per excision event (*attB*), indicating that Tn*Smu1* is capable of autonomous replication, albeit at low copy number. By analogy to ICE*Bs1* and Tn*916*, Tn*Smu1* most likely undergoes rolling-circle replication that initiates from the origin of transfer (*oriT*) using the element-encoded relaxase (NicK).

As the typical lifecycle of *S. mutans* is within biofilms and on the solid surface of teeth, we sought to determine if Tn*Smu1* was capable of excision when cells were grown on a solid surface. *S. mutans* cells were grown to mid-exponential phase, pelleted, and resuspended at a low cell density. Cells were then spotted onto solid medium for 1, 3, 5, or 7 days in anaerobic conditions. We found that excision increased ∼10-fold on solid surfaces compared to liquid culture (Fig. 4B). This likely indicates that Tn*Smu1* is activated when cells are grown on a solid surface, perhaps analogous to the growth in plaque on teeth. This increased activation could also be due to some additional stress experienced during growth on solid surfaces, perhaps resulting in an increase in DNA damage and thus increased activation.

We anticipate that the mechanism of Tn*Smu1* de-repression is analogous to that for ICE*Bs1*. In the presence of DNA damage-inducing conditions, the ICE*Bs1*-encoded metalloprotease ImmA_ICE*Bs1*_ cleaves the repressor ImmR _ICE*Bs1*_, thereby causing de-repression of element gene expression (Auchtung et al., 2007; Bose and Grossman, 2011). Based on the similarities between these two ICE*Bs1* and Tn*Smu1* regulators, the simplest model is that during DNA damage, ImmA_Tn*Smu1*_ becomes activated to cleave and inactivate ImmR_Tn*Smu1*_, thereby causing de-repression of Tn*Smu1* gene expression. We also note that unlike that for ICE*Bs1* (Auchtung et al., 2007; Bose and Grossman, 2011), there are no indications that Tn*Smu1* is regulated by population density, peptide signaling, or quorum sensing.

### Tn*Smu1* can transfer to *S. mutans* recipients that lack a copy of the element

Based on the presence of an apparently intact set of genes for a T4SS and the ability of Tn*Smu1* to excise, we tested for the ability of the element to transfer from one cell to another. To monitor transfer of Tn*Smu1*, we constructed donor and recipient cells that could be detected by their unique antibiotic resistances and auxotrophies.

#### Donors

We introduced a gene conferring tetracycline resistance (*tet*) into Tn*Smu1* between *smu_210c* and *smu_211c* (Fig. 1), generating Tn*Smu1*-*tet* (strain LKM145). This insertion is in a region of Tn*Smu1* with unknown function and we anticipated would not interfere with the typical ICE lifecycle. We found that excision of Tn*Smu1-tet* (∼0.002%, ∼2×10^-5^ *attB/ ilvB*) was similar to that of wild type Tn*Smu1* (Fig. 4A).

We also made a mutant of *S. mutans* that requires D-alanine for growth due to a null mutation in *alr* (*alr*::*erm*), encoding alanine racemase needed for production of D-alanine (Wecke et al., 1997)). Use of an *alr* mutant as a donor enables counter-selection (preventing donor growth) in the absence of D-alanine, analogous to approaches used for *B. subtilis* (Brophy et al., 2018). Donors were also defective in genetic competence (Δ*comS*::*kan*) (Mashburn-Warren et al., 2010) to prevent transformation with DNA from recipients.

#### Recipients

We thought it important to use a recipient that was cured of Tn*Smu1* (Tn*Smu1*^0^). Some ICEs and temperate phages have immunity and exclusion mechanisms that reduce acquisition of additional copies of the cognate element (Auchtung et al., 2007; Gottesman and Weisberg, 2004; Oppenheim et al., 2005; Serfiotis-Mitsa et al., 2008). Notably, ICE*Bs1* has repressor-mediated immunity (Auchtung et al., 2007). Because Tn*Smu1* encodes a homolog of ImmR_ICE*Bs1*_, we were concerned that cells containing Tn*Smu1* might also have repressor-mediated immunity. Therefore, we used two different recipient strains, one without (ΔTn*Smu1*) and one with Tn*Smu1* (LMK85 and LMK87, respectively). Of note, we found there was no noticeable growth difference in strains with or without Tn*Smu1* (Fig. S2). Further, both recipients contained a deletion of *comS* (Mashburn-Warren et al., 2010) to ensure any DNA transfer detected was not via transformation into the recipients.

We found that Tn*Smu1* can transfer from *S. mutans* to *S. mutans,* providing the recipients do not contain a copy of Tn*Smu1. S. mutans* donor and recipient cells were grown to mid-exponential phase, pelleted, and resuspended at a low cell density. Cells were then combined at different starting ratios of donor to recipient cells (1:100, 1:10, 1:1, 10:1, 100:1). Each mixture was spotted onto solid medium for 1, 3, 5, or 7 days. Cells were then harvested and the number of donors, recipients, and transconjugants were enumerated.

We found that Tn*Smu1* transferred from donors into recipients that lacked Tn*Smu1* (Fig. 5 and Fig. S3). Transfer was detected at all donor to recipient ratios and at each of the time points tested (1, 3, 5, 7 days). The largest number of Tn*Smu1* transconjugants were detected at 3 days post-inoculation of the mating plates at a donor to recipient ratio of 1:1. Transfer at this time resulted in ∼10^4^ transconjugants (Fig. 5 and Fig. S3), corresponding to a mating frequency of ∼2 x 10^-5^ when normalized to the number of donors post-mating. The number of transconjugants increased between 1 and 3 days post-inoculation and then dropped at later times (Fig. 5). This increase is due either to transconjugants dividing and producing progeny or transconjugants becoming donors and further transferring Tn*Smu1* to additional recipients. The drop in the number of transconjugants detected at later times was most likely due to overall cell death of donors, recipients, and transconjugants that occurred throughout the mating (Fig. 5B).

**Figure 5.**
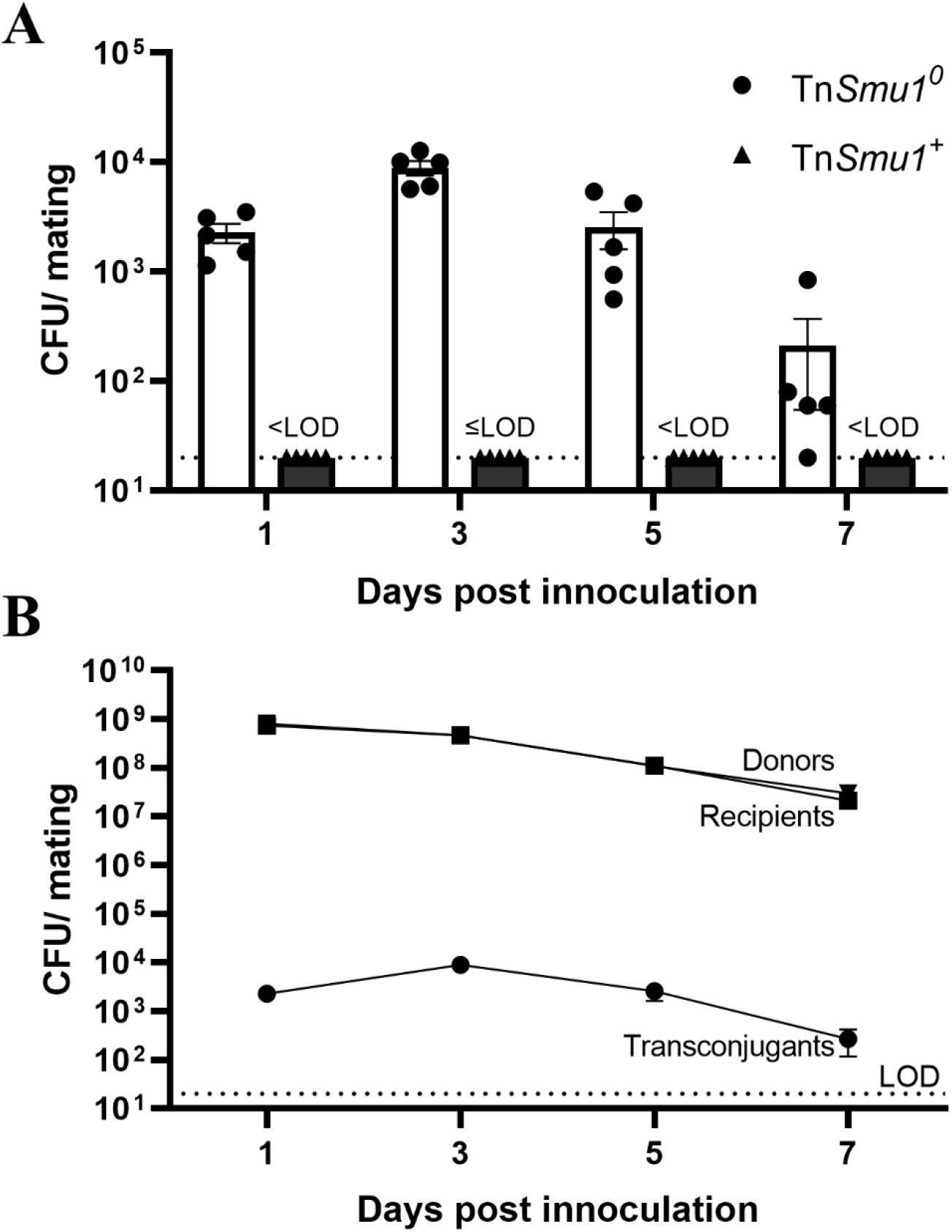
Tn*Smu1* is capable to transfer to recipient cells lacking a copy of the element. Tn*Smu1-tet* donors (LKM145) and Tn*Smu1*^0^ (LKM85) or Tn*Smu1*+ (LKM87) recipients were grown to mid-exponential phase in liquid medium. Donor and recipient cells were pelleted, resuspended at a low density, and mixed at a 1:1 ratio. These mating mixes were spotted onto solid medium for 1, 3, 5, or 7 days in anaerobic conditions. Cells were then harvested and the numbers of donors (*tet*, *alr*, *kan*), recipients (*alr*+, *spc*), and transconjugants (*tet*, *alr+*, *spc*) were enumerated based on unique phenotypes associated with each cell type. **A**. The number of transconjugants formed when Tn*Smu1-tet* donors were co-cultured with recipients without Tn*Smu1*^0^ (LKM85, circles, white bars) or with Tn*Smu1*+ (LKM87, triangles, dark gray bars). Results with Tn*Smu1*^+^ recipients were at or below the limit of detection (LOD ≤20 CFU/mating; dotted line). In some experiments, there was one colony detected. These were not experimentally validated as transconjugants and we indicate this as at the limit of detection (20 CFU/mating). **B.** The numbers of CFUs for donors (inverted triangles), recipients (squares) and transconjugants (circles) in a mating mix are shown for the experiment in Fig. 5A with recipients lacking Tn*Smu1*. The number of transconjugants increased between 1 and 3 days post inoculation and then dropped. The drop in the number of transconjugants at later times was most likely due to overall cell death that occurred throughout the mating as seen with a parallel drop in numbers of donors and recipients. Data presented are averages from three or more independent experiments. Error bars represent the standard error of the mean and are generally not visible as they are too small relative to the size of each data point. Donor and recipient CFUs are largely indistinguishable in this graph. The dotted line at the bottom represents the limit of detection for all cell types.

In contrast to the results with Tn*Smu1*-cured recipients, we detected few if any transconjugants in matings with recipients that contained Tn*Smu1* (Fig. 5A and Fig. S3). Transconjugants were rarely detected, and when they were, we only observed a single colony. We did not verify that these colonies were actual transconjugants, but assuming that they were, then there were ≤20 total transconjugants per mating (at or below the limit of detection) at all time points and donor to recipient ratios tested. This represents a decrease of at least ∼500-fold compared to isogenic recipients without Tn*Smu1*. Together, these results show that Tn*Smu1* is a functional conjugative element, capable of transfer from donor to recipient cells. They also indicate that there is at least one mechanism conferred by Tn*Smu1* that inhibits acquisition of another copy of the element.

### Expression of the repressor, ImmR_Tn_*_Smu1_*, is sufficient to reduce acquisition of Tn*Smu1*

Because Tn*Smu1* encodes a homolog of the immunity repressor ImmR from ICE*Bs1*, we postulated that ImmR_Tn*Smu1*_ might also confer some level of immunity that inhibits acquisition of a second copy of the element, analogous to that of ICEB*s1* (Auchtung et al., 2007; Bose et al., 2008). Therefore, we sought to determine if expression of the repressor encoded by Tn*Smu1*, in the absence of other Tn*Smu1* genes, would result in inhibition of (immunity to) acquisition of Tn*Smu1*.

We found that ectopic expression of ImmR_Tn*Smu1*_ in recipient cells inhibited acquisition of Tn*Smu1*. We used recipients that were missing Tn*Smu1* but expressed *immR*_TnS*mu1*_ under its predicted endogenous promoter and compared these to recipients with and without Tn*Smu1*, analogous to the experiments described above. We measured acquisition of Tn*Smu1* after 3 days of mating at a donor to recipient ratio of 1:1. We found that expression of *immR*_TnS*mu1*_, in the absence of other genes from the element, reduced acquisition of Tn*Smu1* (Fig. 6). This reduction was not as much as that observed with recipients that contained the intact element (Fig. 6). This difference could indicate that there are higher levels of ImmR_TnS*mu1*_ in cells with an intact element, or that there are additional element-encoded mechanisms that contribute to inhibition of a second copy of the element. Together, these results indicate that expression of the Tn*Smu1* repressor (ImmR) is sufficient to inhibit acquisition of the element, and that Tn*Smu1* may have at least one other mechanism that also inhibits acquisition of another copy.

**Figure 6.**
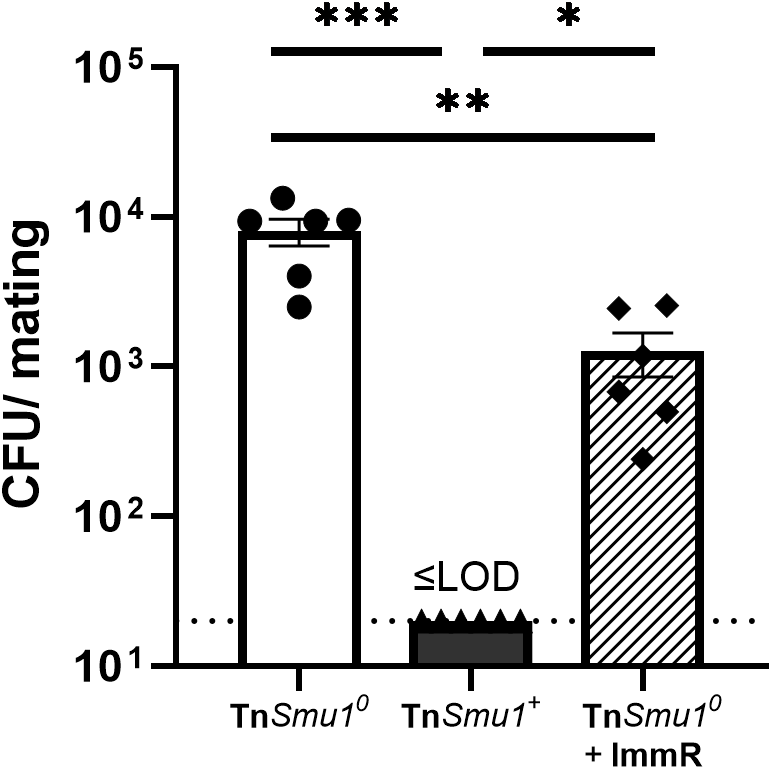
Expression of ImmR_Tn_*_Smu1_* in recipients inhibits acquisition of Tn*Smu1*. Tn*Smu1-tet* donors (LKM145) were mated with three different recipients: Tn*Smu1*^0^ (LKM85, circles, white bar); Tn*Smu1*+ (LKM87, triangles, dark gray bar); and Tn*Smu1*^0^ that expressed *immR*_Tn*Smu1*_ from an ectopic site (LKM178, diamonds, bar with diagonal stripes). Donors and recipients were grown and prepared as described for Fig. 5, except growth on the solid surface (mating) was for 3 days. Data presented are averages from three independent experiments with error bars depicting standard error of the mean. Results with Tn*Smu1*^+^ recipients were at or below LOD (≤20 CFU/mating; dotted line). * p<0.05, **p<0.01, ***p<0.001.

### The preferred integration site of Tn*Smu1* is in a leucyl-tRNA gene

Tn*Smu1* resides in donor cells at the 3’ end of a leucyl tRNA gene (*smu_t33*) (Fig. 3B). We sought to determine if Tn*Smu1* integrated at this same location following transfer to new cells, that is, in transconjugants, or if it was more promiscuous in site selection, perhaps analogous to Tn*916* (Wozniak and Waldor, 2010).

We found that following conjugation, Tn*Smu1* integrated specifically in the 3’ end of *smu_t33*. After performing matings into ΔTn*Smu1* recipient cells (LKM85) as described above, we isolated 16 independent transconjugants and tested these for integration into *smu_t33* using PCR primers that would detect *attL* (Fig. 3A; **Methods**). We found that Tn*Smu1* had integrated into *smu_t33* in all 16 of the transconjugants tested. This tRNA gene (*smu_t33*) and the 17 bp *attB* are not found anywhere else in the *S. mutans* genome (Ajdic et al., 2002). Together, our results indicate that the identified 17 bp site referred to as *attB* (Fig. 3B) is the preferred site of integration of Tn*Smu1*. That said, we cannot rule out the possibility that Tn*Smu1* might integrate into other sites in the chromosome at a low frequency.

### Tn*Smu1* causes a growth arrest in host cells

Tn*916*, an ICE related to Tn*Smu1,* causes growth arrest and death of host cells following excision and expression of its genes (Bean et al., 2022a). In contrast, ICE*Bs1,* which encodes conjugation machinery and regulatory proteins similar to those encoded by Tn*Smu1,* does not cause death of its host cells (Babic et al., 2011; Bean et al., 2022a). Therefore, we wondered whether or not Tn*Smu1* manipulated the growth and-or viability of its host cells. As Tn*Smu1* only excises in a relatively small fraction of cells (Fig. 4), we decided to examine Tn*Smu1* activation in single cells using fluorescence microscopy and a fluorescent reporter that would be indicative of Tn*Smu1* gene expression. We inserted *gfpmut2* between *smu_210* and *smu_211* in Tn*Smu1* (Fig. 1) to generate Tn*Smu1*-*gfp* (LKM137). Cells should fluoresce green only when Tn*Smu1* genes are expressed. Insertion of *gfp* within Tn*Smu1* did not have a significant impact on excision as Tn*Smu1*-*gfp* excised at a frequency similar to that of wild type Tn*Smu1* (∼0.004%, ∼4×10^-5^ *attB/ ilvB*) in liquid medium (Fig. 4A).

We found that cells expressing Tn*Smu1*-*gfp* had a growth defect. Cells containing Tn*Smu1-gfp* were diluted to early exponential phase, grown for three hours, and then spotted onto an agarose pad on a microscope slide. Cells were visualized and tracked for three hours, comparing those that had activated Tn*Smu1-gfp* to those that had not. We tracked 82 cells in which Tn*Smu1-gfp* was activated (GFP on) and 82 neighboring cells in which Tn*Smu1-gfp* was apparently not activated (GFP off) (Fig. 7). Of the cells activating Tn*Smu1* (GFP on), 94% (77/82) did not undergo any further cell divisions and 6% (5/82) divided once (Fig. 7). In contrast, in the 82 neighboring cells without Tn*Smu1*-GFP activated (GFP off), only 4% (3/82) of cells did not undergo any further cell divisions, and 96% of cells underwent 1 or more cell divisions (79/82) (77/82 v. 3/82, χ^2^= 133.64, p<0.0001).

**Figure 7.**
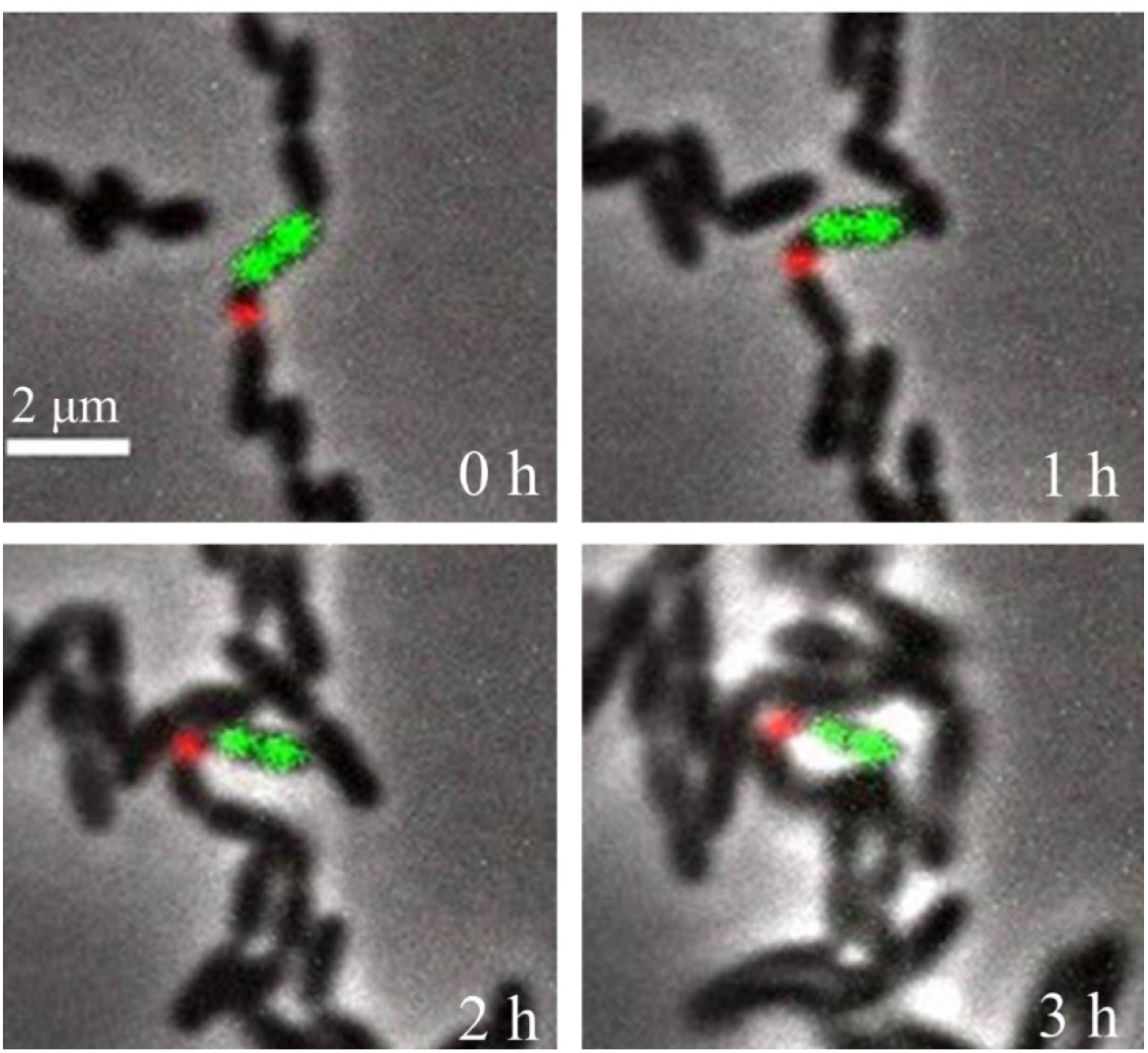
Cells with an active Tn*Smu1* stop growing. Cells containing Tn*Smu1-gfp-tet* (LKM137) were grown to late exponential phase. At time 0 h, cells were spotted on agarose pads containing TH medium, 0.1M propidium iodide, and 0.35 µg/ml DAPI. Cells were monitored by phase contrast and fluorescence microscopy for three hours. GFP (green) was produced in cells in which Tn*Smu1* was activate and excised from the chromosome. Propidium iodide (red) indicates cell death. Images shown are a merge of phase contrast and fluorescence. Three independent experiments gave similar results and were quantified (described in Results). Of 82 cells observed with an active Tn*Smu1-gfp* (GFP on, green) 94% (77/82) did not undergo any further cell divisions and 6% (5/82) divided once. Of 82 neighboring cells that had not activated Tn*Smu1*-*gfp* (GFP off), only 4% (3/82) of cells did not undergo any further cell divisions, and 96% (79/82) of cells underwent one or more cell divisions. A representative set of images is shown here. DAPI is not shown for visual clarity.

Although the cells expressing Tn*Smu1* had a growth arrest, we found no evidence of cell death. Using propidium iodide (PI) to monitor cell viability, we found that of the 82 cells that had activated Tn*Smu1*-GFP, only 12% (10/82) became PI-positive during the course of the experiment (Fig. 7). This was not significantly different than that of cells without Tn*Smu1* activated, where 13% of cells became PI-positive or had progeny that became PI-positive (11/82) throughout the experiment. These numbers are consistent with *S. mutans* viability of randomly selected wild type cells (UA159, 10/82 cells became PI-positive or had progeny that became PI-positive over the 3 hours of the experiment) and cells without Tn*Smu1* (LKM68, 9/82 cells became PI-positive or had progeny that became PI-positive over the 3 hours of the experiment). This is also consistent with what is seen in *S. mutans* biofilms and various growth conditions via fluorescence microscopy (Decker, 2001; Decker et al., 2014; Zhang et al., 2009). Together, these results indicate that cells in which Tn*Smu1* becomes activated are unable to divide; however, they do not lose viability, at least over a period of three hours.

## Discussion

Our work demonstrates that Tn*Smu1* is a functional ICE, capable of excision from the host cell chromosome and transfer to recipient cells. We found that activation of Tn*Smu1* is stimulated by DNA damage and by growth on a solid surface. Further, activation of Tn*Smu1* caused a growth arrest, indicating that it can have a profound effect on host physiology. Our findings provide a basis for future work examining the impacts of Tn*Smu1* on the physiology of *S. mutans*. Further, as ICEs can be utilized for genetic engineering purposes (Bean et al., 2022b; Brophy et al., 2018; Miyazaki and van der Meer, 2013; Peters et al., 2019), Tn*Smu1* may perhaps be developed as an engineering tool for manipulation of organisms in the oral microbiome.

### The Tn*Smu1* repressor

*immR*_Tn*Smu1*_ (*smu_218c*) almost certainly encodes the repressor of Tn*Smu1*, based on three lines of evidence. First, ImmR_Tn*Smu1*_ is similar to the repressor of ICE*Bs1* and others in this family. Second, like other ‘immunity’ repressors, expression in recipient cells (in the absence of other element genes) caused a reduction in acquisition of a copy of the element, similar to immunity to superinfection exhibited by various temperate phages (Gottesman and Weisberg, 2004; Oppenheim et al., 2005) and ICE*Bs1* (Auchtung et al., 2007; Bose et al., 2008). Lastly, *smu_218c* (*immR*) appears to be an essential gene based on Tn-seq experiments with *S. mutans* (Shields et al., 2018). Repressors of mobile genetic elements that integrate and excise (e.g., ICEs and temperate phages) can appear to be ‘essential’ because their loss can result in excision and loss of the element in which they are encoded. For example, if loss of a repressor leads to excision of the element, that element will likely be lost from the population of cells. This loss makes it very difficult to establish a null mutation in the gene for the repressor as the loss of function mutation in the repressor gene will be lost along with the resulting excised element. In this way, genes potentially encoding element repressors can be identified in genome-wide screens and appear as ‘essential’ even though they reside in an element that itself is not essential. This point is highlighted by the fact that we were able to delete Tn*Smu1* and cells are viable, but the repressor appears to be essential (Shields et al., 2018).

### Costs and benefits to cells carrying an ICE

ICEs are often double-edged swords to their host cells: They can provide both fitness costs and benefits. They can benefit their host cells through associated cargo genes that confer specific phenotypes, such as antibiotic resistances, metabolic traits, and virulence factors. However, some ICEs can manipulate host development, growth, and viability (Beaber et al., 2004; Bean et al., 2022a; Jones et al., 2021; Pembroke and Stevens, 1984; Reinhard et al., 2013). This is similar to plasmids, which are known to provide beneficial phenotypes to their hosts but are energetically costly to maintain (San Millan and MacLean, 2017).

This complex interplay between element and host is evident in Tn*916* and *Pseudomonas* ICE*clc*. Tn*916* was discovered through its ability to spread tetracycline resistance through clinical isolates of *Enterococcus* (Franke and Clewell, 1981a, 1981b), thus providing a clear benefit to its host cells. However, activation of Tn*916* halts cell growth and leads to decreased bacterial viability (Bean et al., 2022a). Similarly, when activated, ICE*clc* causes slow growth and decreased viability (Reinhard et al., 2013). These growth defective cells are in a “transfer competent state”. Deletion of the genes required for the decreased cell growth and viability cause a decrease in conjugation efficiency. This indicates that this state of decreased cell growth and viability is important for efficient transfer of ICE*clc* (Delavat et al., 2016; Reinhard et al., 2013).

There are certainly parallels between the growth arrest caused by Tn*Smu1,* Tn*916,* and ICE*clc*. However, unlike Tn*916* and ICE*clc*, our results indicate that Tn*Smu1* does not cause host cell death. Additionally, a CRISPRi knockdown of *immR*_Tn*Smu1*_ caused an arrest in growth of the entire population (King et al., 2022). It is possible that the cells with an activated Tn*Smu1* die following growth arrest, but we have not observed this, nor have assays measuring death been reported. The apparent essentiality of *immR*_Tn*Smu1*_ (Shields et al., 2018) could indicate that there is cell death following inactivation of the repressor and subsequent activation of Tn*Smu1*. However, as discussed above, we postulate that this apparent essentiality is due to excision and loss of Tn*Smu1* and the consequent loss of the *immR* null allele in Tn*Smu1*. This further begs the questions: What is the mechanism of the growth arrest caused by activation of Tn*Smu1*, is this growth arrest important for conjugation and transfer and is there a benefit that Tn*Smu1* confers to host cells.

### Host range of Tn*Smu1*

Bioinformatic searches indicate that Tn*Smu1* is naturally located within *S. mutans* species. However, it is not known if Tn*Smu1* is capable of transfer to other bacterial species typically found in the oral cavity. ICE*Bs1* is naturally only found in *Bacillus* sp. but can transfer to a diverse array of other microbes (Auchtung et al., 2005; Brophy et al., 2018). Tn*916* is naturally found in *Enterococcus*, *Clostridium*, *Streptococcus*, and *Staphylococcus* species, and is also functional in *Bacillus* sp. (Roberts and Mullany, 2009, 2011; Wright and Grossman, 2016). Further, many ICEs are able to mediate transfer of other mobile genetic elements, including plasmids that replicate by rolling-circle replication (Johnson and Grossman, 2015; Lee et al., 2012; Santoro et al., 2014). We suspect that Tn*Smu1* is able to drive transfer of plasmids and other mobile genetic elements found within the oral microbiome. As many mobile genetic elements encode antibiotic resistance, the impact of Tn*Smu1* on clinically important phenotypes (virulence traits and drug resistances) may extend beyond transfer of Tn*Smu1* alone.

## Materials & Methods

### Media and growth conditions

For liquid growth, *S. mutans* cultures were grown statically in 50% Todd Hewitt (TH) broth in tightly closed 15 ml conical tubes. For growth on solid media, *S. mutans* were grown on Brain Heart Infusion plates with 1.5% agar under anaerobic conditions (using Anaerogen Anaerobic Gas Generator, Hardy Diagnostics). All growth occurred at 37°C. When appropriate, media was supplemented with 1.6 mg/ml D-alanine. Antibiotics were used at the following concentrations: 1 mg/ml kanamycin, 1 mg/ml spectinomycin, 10 μg/ml erythromycin.

### Strains and alleles

*S. mutans* strains (Table 2) were derived from *S. mutans* UA159 (ATCC 700610) (Ajdic et al., 2002) and were made by natural transformation (Li et al., 2001; Petersen and Scheie, 2010). New alleles were constructed as double crossover events using long-flanking homology PCR by isothermal assembly (Gibson, 2009; Xie et al., 2011). Markers used to select for transformants were *spc* (spectinomycin resistance), *erm* (erythromycin resistance), *tet* (tetracycline resistance) or *kan* (kanamycin resistance). All mutants were constructed in a clean, isogenic background and alleles were confirmed through Sanger sequencing. Any alleles obtained from other sources were moved into a clean isogenic background. Construction of new stains and alleles is summarized below.

**Table 2.**
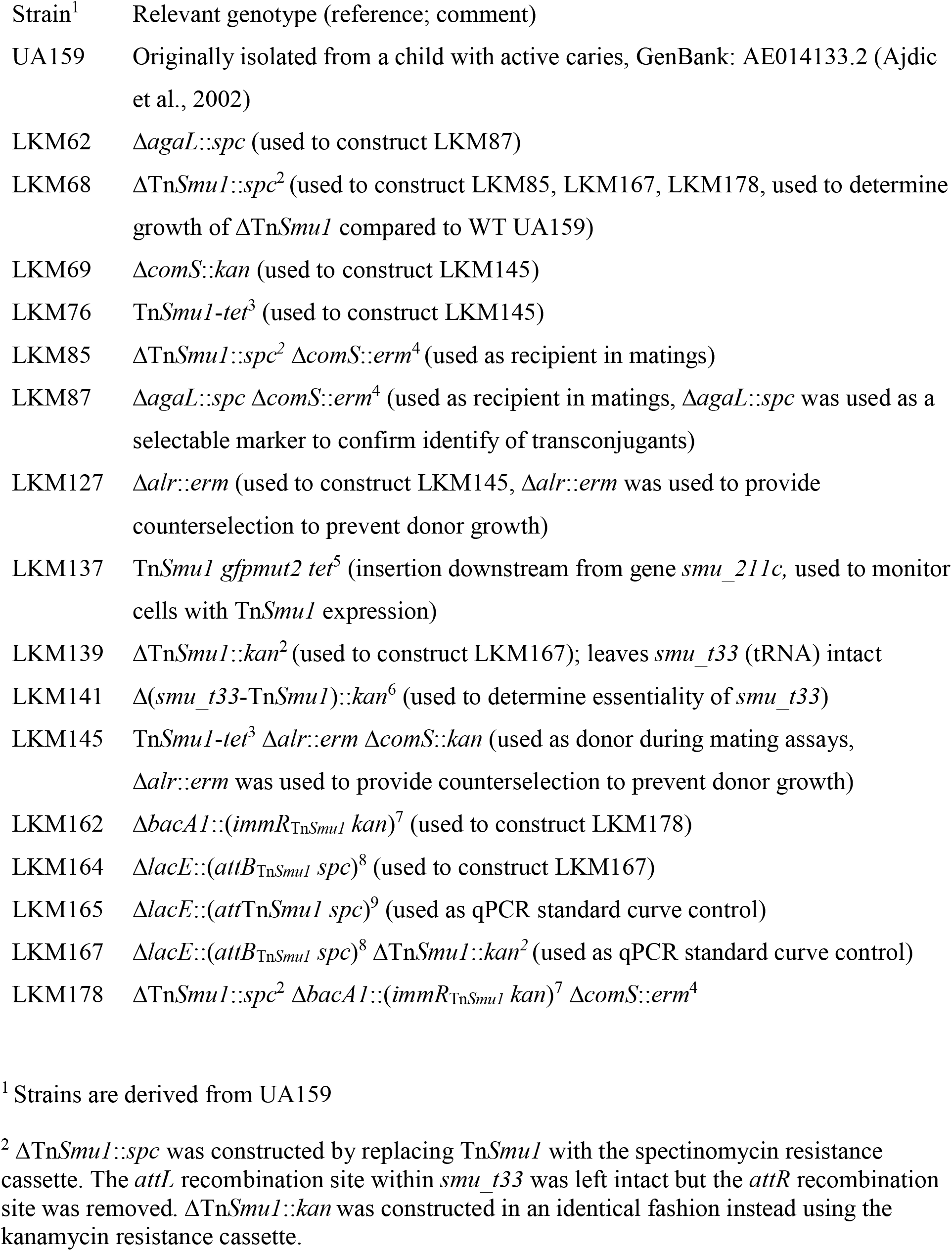
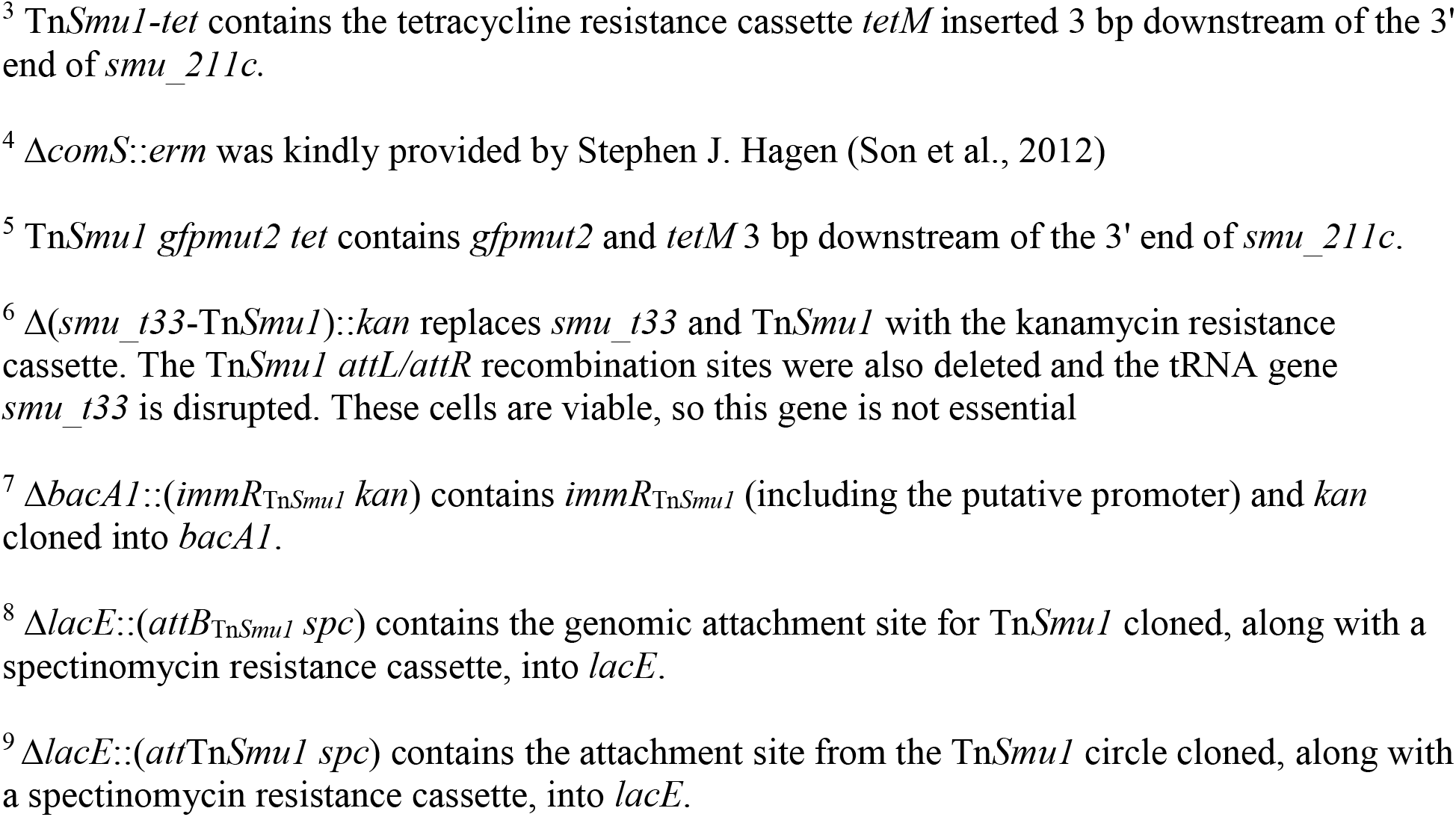
*S. mutans* strains.

| Strain<sup>1</sup> | Relevant genotype (reference; comment) |
| --- | --- |
| UA159 | Originally isolated from a child with active caries, GenBank: AE014133.2 (Ajdic et al., 2002) |
| LKM62 | $\Delta agaL::spc$ (used to construct LKM87) |
| LKM68 | $\Delta TnSmuI::spc^2$ (used to construct LKM85, LKM167, LKM178, used to determine growth of $\Delta TnSmuI$ compared to WT UA159) |
| LKM69 | $\Delta comS::kan$ (used to construct LKM145) |
| LKM76 | $TnSmuI-tet^3$ (used to construct LKM145) |
| LKM85 | $\Delta TnSmuI::spc^2 \Delta comS::erm^4$ (used as recipient in matings) |
| LKM87 | $\Delta agaL::spc \Delta comS::erm^4$ (used as recipient in matings, $\Delta agaL::spc$ was used as a selectable marker to confirm identify of transconjugants) |
| LKM127 | $\Delta alr::erm$ (used to construct LKM145, $\Delta alr::erm$ was used to provide counterselection to prevent donor growth) |
| LKM137 | $TnSmuI gfpmut2 tet^5$ (insertion downstream from gene *smu_211c*, used to monitor cells with *TnSmuI* expression) |
| LKM139 | $\Delta TnSmuI::kan^2$ (used to construct LKM167); leaves *smu_t33* (tRNA) intact |
| LKM141 | $\Delta (smu_t33-TnSmuI)::kan^6$ (used to determine essentiality of *smu_t33*) |
| LKM145 | $TnSmuI-tet^3 \Delta alr::erm \Delta comS::kan$ (used as donor during mating assays, $\Delta alr::erm$ was used to provide counterselection to prevent donor growth) |
| LKM162 | $\Delta bacAI::(immR_{TnSmuI} kan)^7$ (used to construct LKM178) |
| LKM164 | $\Delta lacE::(attB_{TnSmuI} spc)^8$ (used to construct LKM167) |
| LKM165 | $\Delta lacE::(attTnSmuI spc)^9$ (used as qPCR standard curve control) |
| LKM167 | $\Delta lacE::(attB_{TnSmuI} spc)^8 \Delta TnSmuI::kan^2$ (used as qPCR standard curve control) |
| LKM178 | $\Delta TnSmuI::spc^2 \Delta bacAI::(immR_{TnSmuI} kan)^7 \Delta comS::erm^4$ |

<sup>1</sup> Strains are derived from UA159
<sup>2</sup> $\Delta TnSmuI::spc$ was constructed by replacing *TnSmuI* with the spectinomycin resistance cassette. The *attL* recombination site within *smu\_t33* was left intact but the *attR* recombination site was removed. $\Delta TnSmuI::kan$ was constructed in an identical fashion instead using the kanamycin resistance cassette.
<sup>3</sup> *TnSmuI-tet* contains the tetracycline resistance cassette *tetM* inserted 3 bp downstream of the 3' end of *smu\_211c*.
<sup>4</sup> $\Delta comS::erm$ was kindly provided by Stephen J. Hagen (Son et al., 2012)
<sup>5</sup> *TnSmuI gfpmut2 tet* contains *gfpmut2* and *tetM* 3 bp downstream of the 3' end of *smu\_211c*.
<sup>6</sup> $\Delta(sm u\_t33-TnSmuI)::kan$ replaces *smu\_t33* and *TnSmuI* with the kanamycin resistance cassette. The *TnSmuI attL/attR* recombination sites were also deleted and the tRNA gene *smu\_t33* is disrupted. These cells are viable, so this gene is not essential
<sup>7</sup> $\Delta bacA1::(immR_{TnSmuI} kan)$ contains *immR<sub>TnSmuI</sub>* (including the putative promoter) and *kan* cloned into *bacA1*.
<sup>8</sup> $\Delta lacE::(attB_{TnSmuI} spc)$ contains the genomic attachment site for *TnSmuI* cloned, along with a spectinomycin resistance cassette, into *lacE*.
<sup>9</sup> $\Delta lacE::(attTnSmuI spc)$ contains the attachment site from the *TnSmuI* circle cloned, along with a spectinomycin resistance cassette, into *lacE*.

ΔTn*Smu1*::*spc* (constructed in LKM68 then transferred via natural transformation to LKM85 and LKM178) was constructed by replacing Tn*Smu1* with the spectinomycin resistance cassette from pUS19 (Benson and Haldenwang, 1993). The *attL* recombination site within *smu_t33* was left intact (leaving 4 bp downstream of the 3’ end of *smu_t33*) but the *attR* recombination site was removed (preserving 67 bp downstream of the 3’ end of *smu_221c*). The allele was constructed via isothermal assembly of the antibiotic cassette and ∼750 bp of upstream and downstream homology arms.

ΔTn*Smu1*::*kan* (constructed in LKM139 then transferred via natural transformation to LKM167) was made in the same way as ΔTn*Smu1*::*spc*, with the same boarders, except that the kanamycin resistance cassette from pGK67 (Lemon et al., 2001) was used instead of *spc*.

Δ(*smu_t33*-Tn*Smu1*)::*kan* in LKM141 deletes Tn*Smu1* and the tRNA gene *smu_t33* and inserts *kan* from pGK67. The deletion endpoints extend from 14 bp downstream of *smu_t32* through *attR,* leaving *smu_221c* (the gene downstream of Tn*Smu1*) intact, and ending 67 bp downstream from the *smu_221c* stop codon. The allele was constructed via isothermal assembly of the antibiotic cassette and ∼750 bp of upstream and downstream homology arms. Although *smu_t33* encodes a unique tRNA, it was not essential: the deletion was viable, albeit with a growth defect.

Δ*agaL*::*spc* (constructed in LKM62 and then transferred via natural transformation to LKM87) was constructed by replacing the *agaL* open reading (maintaining the first 558 bp and last 458 bp of the *agaL* open reading frame) with the spectinomycin resistance cassette from pUS19. The allele was constructed via isothermal assembly of the antibiotic cassette and ∼750 bp of upstream and downstream homology arms. *agaL* is considered a non-essential chromosomal location suitable for cloning (Reck and Wagner-Döbler, 2016).

Tn*Smu1-tet* (initially in LKM76 and then transferred via natural transformation to make LKM145) was constructed by inserting the tetracycline resistance cassette (*tetM*) 3 bp downstream of the 3’ end of *smu_211c*. The allele was constructed via isothermal assembly of the antibiotic cassette and ∼750 bp of upstream and downstream homology arms. *tetM* from Tn*916* was used to confer tetracycline resistance from CMJ253 (including 376 bp upstream of *tetM* so as to include the *tetM* promoter) (Johnson and Grossman, 2014). *tetM* was co-directional with the upstream and downstream genes in Tn*Smu1* and did not contain a transcriptional terminator downstream of *tetM*. Tn*Smu1 gfpmut2-tet* in LKM137 was constructed in an identical manner by inserting *gfpmut2* allele and *tetM* from Tn*916* from CMJ253 (Johnson and Grossman, 2014) 3 bp downstream of the 3’ end of *smu_211c*. *gfpmut2* was obtained from ELC1458 (Bean et al., 2022a). *gfpmut2* is promoter-less and co-directional with the upstream and downstream genes in Tn*Smu1.* It has the *B. subtilis spoVG* ribosome binding site to initiate translation.

*Δalr:*:*erm* (initially in LKM127 and then transferred via natural transformation to make LKM145) was constructed via isothermal assembly of the erythromycin antibiotic cassette (*erm*) and ∼750 bp of upstream and downstream homology arms. The first 4 bp of the *alr* open reading frame was retained as it overlapped with *acpS* but the rest of the *alr* open reading frame was deleted (to 172 bp upstream of the 5’ end of *recG*). The erythromycin resistance cassette from pCAL215 was used (Auchtung et al., 2007), originally derived from pDG795 (Guérout-Fleury et al., 1996).

Δ*comS*::*kan* (originally in LKM69 and then transferred via natural transformation to make LKM145) was constructed by replacing the entire *comS* open reading frame with the kanamycin resistance cassette from pGK67. The allele was constructed via isothermal assembly of the antibiotic cassette and ∼750 bp of upstream and downstream homology arms. The Δ*comS*::*erm* allele was kindly provided by Stephen J. Hagen and construction was previously described (Son et al., 2012). This allele was transferred via natural transformation to LKM85, LKM87, and LKM178.

Δ*bacA1*::(*immR*_Tn*Smu1*_ *kan*) (*smu218c kan*) (initially in LKM162 and then transferred via natural transformation to make LKM178) was constructed by replacing the *bacA1* open reading frame with *immR*_Tn*Smu1*_ and *kan* (maintaining the first 511 bp and last 548 bp of the *bacA1* open reading frame)*. immR*_Tn*Smu1*_ was amplified from Tn*Smu1* from S*. mutans* UA159, amplifying 746 bp upstream of the 5’ end of *immR*_Tn*Smu1*_ such that it contains the predicted promoter driving expression of *immR*_Tn*Smu1*_, to the end of the *immR*_Tn*Smu1*_ open reading frame. The organization of Tn*Smu1* is reminiscent of that of ICE*Bs1* where *immR* and *immA* are co-transcribed and divergent from most of the genes in the element (Auchtung et al., 2007; Bose et al., 2008). Therefore, we predicted the intergenic region between *smu_227* and *immR*_TnS*mu1*_ (*smu_228*) would contain the promoter of ImmR_Tn*Smu1*_. The kanamycin resistance cassette from pGK67 was cloned divergent to *immR*_Tn*Smu1*_. The allele was constructed via isothermal assembly of the antibiotic cassette, the *immR*_Tn*Smu1*_ fragment, and ∼750 bp of upstream and downstream homology arms. *bacA1* is a non-essential chromosomal location suitable for cloning (Reck and Wagner-Döbler, 2016).

Δ*lacE*::(*att*Tn*Smu1 spc*) in LKM165 was constructed by inserting *att*Tn*Smu1* and *spc* into *lacE*, maintaining the first 512 bp and the last 463 bp of the *lacE* open reading frame. *att*Tn*Smu1* and the surrounding regions in Tn*Smu1* (containing the last 730 bp of the *int*_Tn*Smu1*_ open reading frame and the first 157 bp of the *smu_220* open reading frame) was amplified from Tn*Smu1* that had spontaneously excised from liquid cultures of *S. mutans* UA159. The spectinomycin resistance cassette from pUS19 was used. The allele was constructed via isothermal assembly of the antibiotic cassette, the *att*Tn*Smu1* fragment, and ∼750 bp of upstream and downstream homology arms. *lacE.* is considered a non-essential chromosomal location suitable for cloning (Reck and Wagner-Döbler, 2016).

Δ*lacE*::(*attB*_Tn*Smu1*_ *spc*) (initially in LKM164 and then transferred via natural transformation to make LKM167) was constructed by inserting *attB*_Tn*Smu1*_ and *spc* into *lacE* (with regions of *lacE* as described above). *attB*_Tn*Smu1*_ was constructed by amplifying the genomic region upstream of *attL* (including the last 16 bp of *smu_t24* to 5 bp downstream of the 3’ end of *smu_*t33 so that the recombination site *attB* is present) and stitching it together with the genomic region downstream *attR* (from 67 bp downstream of the 3’ end of *smu_221* to 124 bp upstream of the 5’ end of *smu_221*) and cloned into pCAL1422 (Thomas et al., 2013). The *attB*_Tn*Smu1*_ was amplified from the resulting plasmid and the spectinomycin resistance cassette was amplified from pUS19. The allele was constructed via isothermal assembly of the antibiotic cassette, the *attB*_Tn*Smu1*_ fragment, and ∼750 bp of upstream and downstream homology arms.

### Homology, bioinformatic analyses, and Tn*Smu1* conservation

Global alignments between Tn*Smu1* were calculated with EMBOSS Needle pairwise global sequence alignment (Needleman and Wunsch, 1970). BLASTp and BLASTn publicly available databases were accessed on June 9, 2022. Structural predictions were done using Phyre2, accessed June 9, 2022. Helix-turn-helix domains were predicted using GYM 2.0 (Gao et al., 1999; Narasimhan et al., 2002).

We used cblaster (Gilchrist et al., 2021) to look at conservation of Tn*Smu1* across publicly available sequences of *S. mutans* accessed most recently June 22, 2022. We used the predicted protein sequences of the genes within Tn*Smu1* as the query for cblaster. Subjects were grouped into clusters based on BLASTp hits to Tn*Smu1* protein sequence and ranked by cluster similarity. Cluster similarity score is calculated by cblaster as *S*=*h*+*i•s*, where *h* is the number of query sequences with BLASTp hits, *s* is the number of contiguous gene pairs with conserved synteny and *i* is a weighting factor (default value 0.5) determining the weight of synteny in the similarity score. If a *S. mutans* strain appeared twice due to multiple copies of the genome sequence available in the NCBI database, the genome with the lower cluster similarity score was excluded from Fig. S1.

### Determination of Tn*Smu1* excision

Genomic DNA was isolated from overnight cultures of *S. mutans* UA159 using Qiagen DNeasy kit with 40 µg/ml lysozyme. Tn*Smu1* excision and the resulting *attB* sequence was identified with primers oLM27 (5’ – ACACCAGATTGTGGCTCTG) and oLM49 (5’ – GGCAAGTCTTGATTATCGCTTTTAGAAAGAG). The *att*Tn*Smu1* junction formed via site-specific recombination was determined using primers oLM38 (5’ – CATCAAGTTAGCACAGTCAGATAAAATCG) and oLM107 (5’ – CATAATAGGTTCCATTTAAACTACTGCC). The resulting products were determined by Sanger sequencing.

### Quantitative PCR to determine excision and replication of the element

Overnight cultures were diluted to OD = 0.05 in 50% TH medium and grown at 37°C. After 3 hours of growth, the culture was split and 1 µg/ml Mitomycin C (MMC) was added to half of the culture (where indicated). Samples were taken every hour pre- and post-MMC addition and into stationary phase (7 hours total). Genomic DNA was isolated using Qiagen DNeasy kit with 40 µg/ml lysozyme. Excision was measured using primers oLM166 (5’ – TTGGTTCGAATCCAGCTACC) and oLM109 (5’-GACTTATGGTCATTTGGTTGCG) to amplify the vacant insertion site *attB. attB* amplification was normalized to a control chromosomal region in *ilvB*, which is ∼7 kb downstream of *attB. ilvB* was amplified with primers oLM173 (5’ –AGGTGGCGGTGTCAATTATG) and oLM174 (5’ – GCATCTCCCACAACTGGAATAG). The copy number of the Tn*Smu1* circle was measured with primer pair oLM224 (5’ –AATCTTCTATCCCAAATTTTCTCCC) and oLM226 (5’ – TGGGAGAAATTTTGGGAGAGAAAATC) to quantitate the unique *att*Tn*Smu1* junction formed via site-specific recombination. To determine if Tn*Smu1* was replicating, we determined the ratio of the number of copies of circular Tn*Smu1* (*att*Tn*Smu1*) to the number of copies of the excision site (*attB*).

qPCR was done using SSoAdvanced SYBR master mix and CFX96 Touch Real-Time PCR system (Bio-Rad). Copy numbers of *attP* and *attB* were determined by the Pfaffl method (Pfaffl, 2001). Standard curves for *att*Tn*Smu1*, *attB*, and the control chromosome locus *ilvB* were generated from genomic DNA of LKM165, LKM167, and *S. mutans* UA159 respectively. LKM165 contains an ectopic copy of *att*Tn*Smu1* inserted at *lacE*. LKM167 does not contain Tn*Smu1* and therefore contains a copy of the unoccupied chromosome site *attB* inserted at *lacE*. *S. mutans* UA159 is wild type *S. mutans* and was used for measuring the nearby chromosomal locus, *ilvB*.

### Growth curves

Strains were grown in 50% TH broth overnight. Cultures were diluted to an OD_600_ of 0.05 and grown statically in closed 15 ml conical tubes. The number of colony forming units (CFUs) and OD_600_ was determined every hour. Cells were monitored for 7 hours of growth.

### Mating assays

Donor and recipient cells were diluted to early exponential phase and grown to mid-exponential phase over 4 hours. Cells were then pelleted at 5000 rpm for 5 mins and resuspended to an OD of 0.01 in 1x Spizizen’s salts (Harwood and Cutting, 1990). Donor and recipient cells were combined at various ratios and 50μl of each mix was spotted onto a BHI plate supplemented with D-alanine. Mating plates were then incubated at 37 °C in anaerobic conditions (using Anaerogen Anaerobic Gas Generator, Hardy Diagnostics) for up to 7 days. Spots were then harvested by scrapping the spot into 1x Spizizen’s salts and vortexed. Cells were then plated on selective media to detect Tn*Smu1* transfer. The number of donor (tetracycline resistant, D-alanine auxotrophs), recipient (tetracycline sensitive, D-alanine phototrophs), and transconjugant (tetracycline resistant, D-alanine phototrophs) CFUs were enumerated both pre- and post-mating. Conjugation efficiency was calculated as the percentage of transconjugant CFUs formed normalized to the number of donor cells harvested at the end of the mating to account for cell growth.

### Time-lapse microscopy and analysis

*S. mutans* cells were diluted to early exponential phase in 50% TH media. After 3 hours of growth to late exponential phase, cells were transferred to an agarose pad (1.5% UltraPure agarose, Invitrogen; dissolved in growth medium) containing 0.1M propidium iodide and 0.35 µg/ml DAPI. The agarose pad was created in an incubation chamber, which was made by stacking two Frame-Seal Slide Chambers (Bio-Rad) on a standard microscope slide (VWR). Cells were then grown at 37°C for 3 hours while monitoring growth. Fluorescence was generated using a Nikon Intensilight mercury illuminator through appropriate sets of excitation and emission filters (Chroma; filter sets 49000, 49002, and 49008). Time-lapse images were captured on a Nikon Ti-E inverted microscope using a CoolSnap HQ camera (Photometrics). ImageJ software was used for image processing and analysis.

## Acknowledgements

We thank Katharina Ribbeck (MIT, Cambridge, MA), Dan Smith (Forsyth Institute, Cambridge, MA) for sharing strains. We thank Stephen J. Hagen (University of Florida, Gainesville, FL) for kindly sharing strains and the Δ*comS*::*erm* allele, James S. Weagley (Washington University, St. Louis, MO) for thoughtful conversions on bioinformatic analyzes of Tn*Smu1* distribution among bacterial species and Sang-Joon Ahn (University of Florida, Gainesville, FL) for useful conversations.

Research reported here is based upon work supported, in part, by the National Institute of General Medical Sciences of the National Institutes of Health under award number R35 GM122538 to ADG. Any opinions, findings, and conclusions or recommendations expressed in this report are those of the authors and do not necessarily reflect the views of the National Institutes of Health.

---

**Figure S1.**
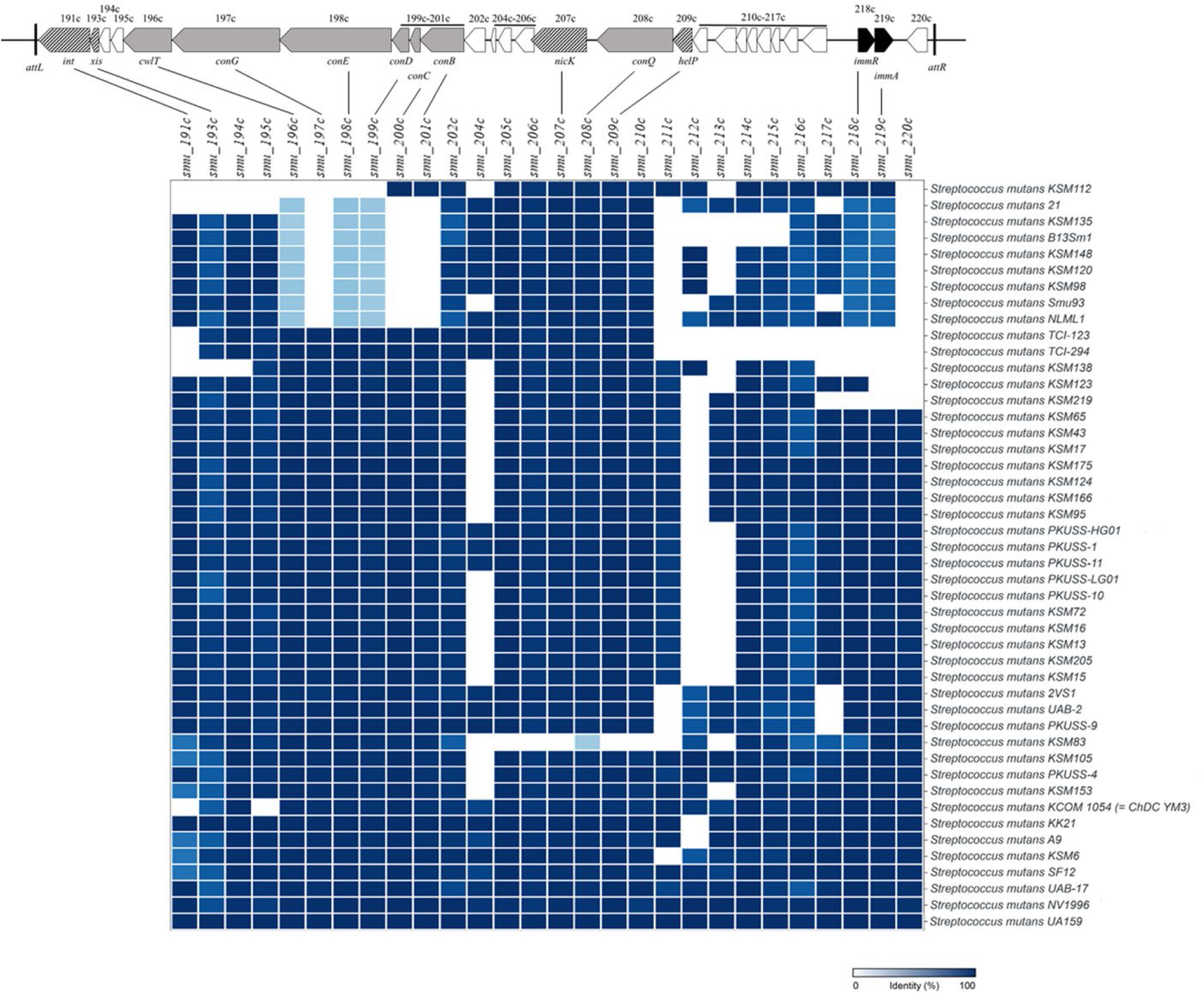
Distribution of Tn*Smu1* among isolates of *S. mutans.* We searched the NCBI BLASTp database for proteins that are similar to those encoded by Tn*Smu1* using cblaster (Gilchrist et al., 2021). Rows represent sequenced *S. mutans* strains and columns represent proteins similar to those encoded by Tn*Smu1.* Boxes represent presence of a protein similar to one encoded by Tn*Smu1,* with the intensity of the blue presenting the percent identity. Sequences were grouped into clusters based on matches with Tn*Smu1* protein sequences and clusters were ranked by similarity. A map of Tn*Smu1* is shown with the open reading frames used for sequence comparisons. Genes names are abbreviated to include only the number designation (e.g., *191c* indicates *smu_191c*). The homologous ICE*Bs1* gene is indicated below the map. Putative gene function is indicated by pattern and color: genes of unknown function (white); type 4 secretion system proteins (gray); DNA processing (diagonal stipes), and regulation (black). We defined Tn*Smu1* as present in a species if the genome contained the genes required for the ICE lifecycle (DNA processing (*smu_191c, smu_193c, smu_207c, smu_209c*), conjugation (*smu_196c-smu_201c, smu_208c)*, and regulation genes (*smu_218c, smu_219c*)). It is possible that elements that contain all the genes needed for a function ICE but not the regulatory genes are functional, but regulated in some other way. Nonetheless, based on the definition including *smu_218c* and *smu219c*), 30 sequenced *S. mutans* genomes contain Tn*Smu1.* Data displayed had a similarity score of >25. *S. mutans* UA159 was used as the query and is shown as a reference.

**Figure S2.**
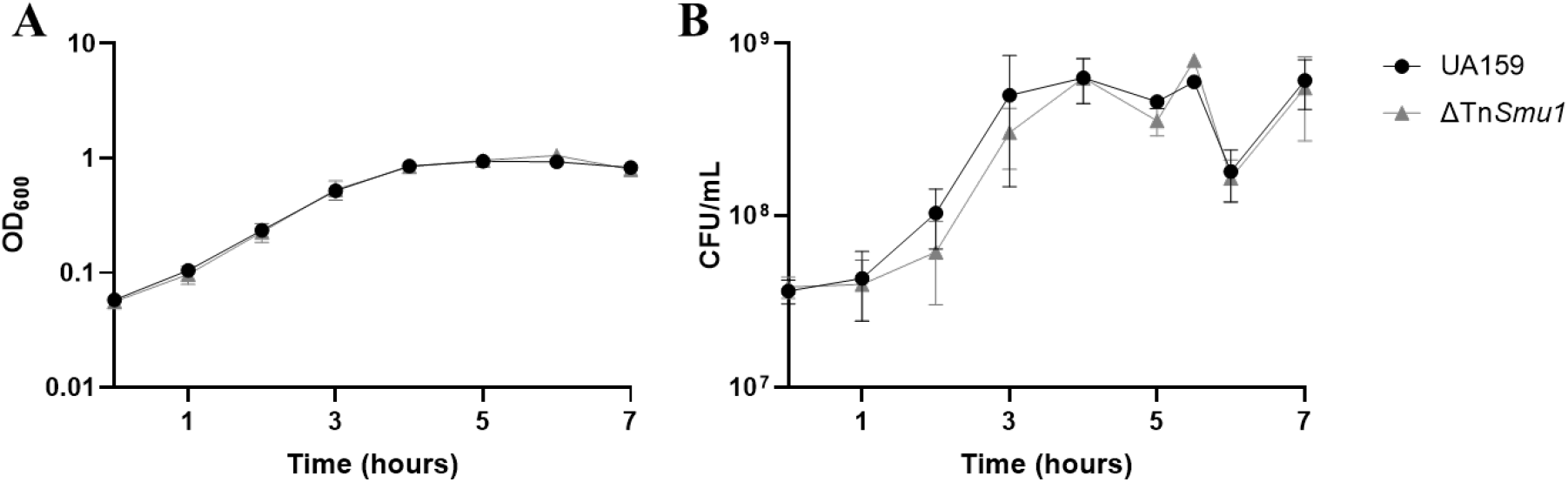
Cells with and without Tn*Smu1* appear to grow similarly. Growth and viability of UA159 (gray circles) and UA159ΔTn*Smu1* (LKM68) (black triangles) was monitored for 7 hours. Optical density (OD_600_; panel A) and CFU/ml (panel B) were measured over time. Data presented are averages from three or more independent experiments with error bars depicting standard error of the mean. For OD measurements (A), those of the two strains largely overlapped and are indistinguishable in the graph and error bars could not be depicted due to the size of each data point.

**Figure S3.**
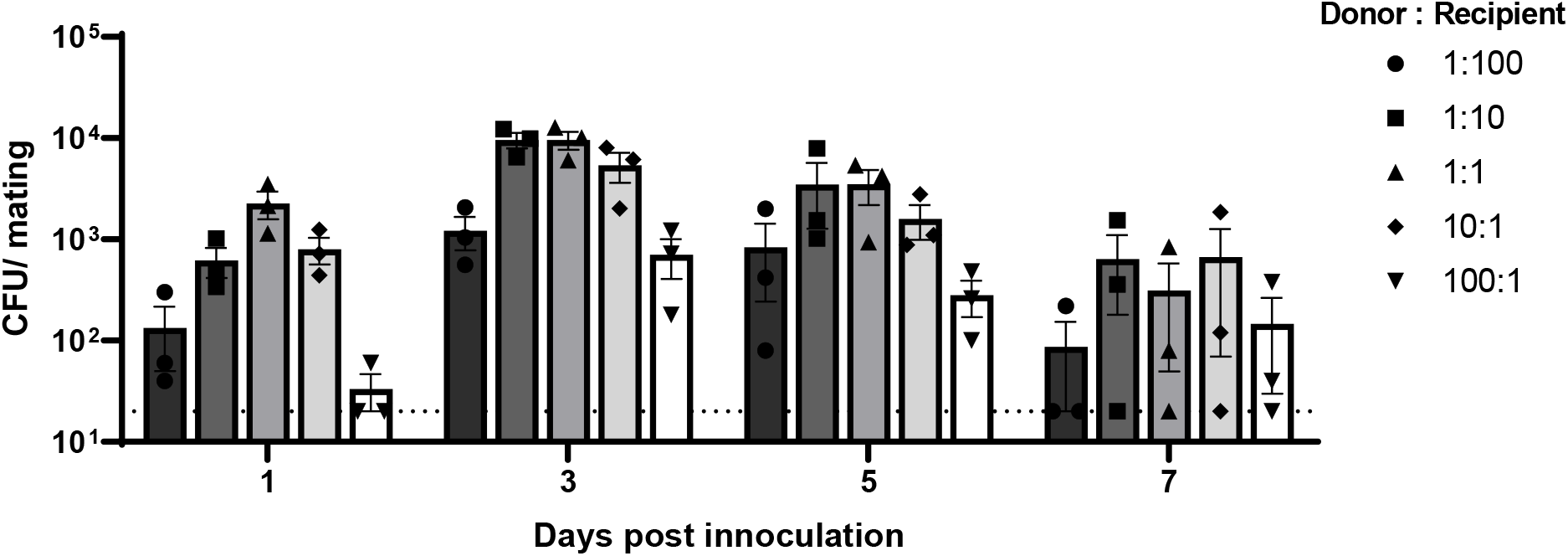
Tn*Smu1* transfer to recipient cells lacking Tn*Smu1.* Tn*Smu1* donors (LKM145) and Tn*Smu1*^0^ (LKM85) recipients were grown to mid-exponential phase over 3 hours. Donor and recipient cells were pelleted and resuspended at a low OD, and mixed at various donor to recipient ratios. Bacteria were then spotted onto solid medium and allowed to grow for 1, 3, 5, or 7 days in anaerobic conditions. Cells were harvested and the numbers of transconjugants were determined. Data are presented from three independent experiments with each bar representing the mean and error bars depicting standard error of the mean. Ratios of donors to recipients included: 1:100, circles and black bars; 1:10, squares and dark gray bars; 1:1, triangles and medium gray bars; 10:1, diamonds and light gray bars; and 100:1, inverse triangles, white bars). Matings using Tn*Smu1*^+^ recipients (LKM87) were done in parallel and the number of transconjugants under all conditions tested was at or below the limit of detection (dotted line; ≤20 CFU/mating).

## Literature cited

Ajdic, D., McShan, W.M., McLaughlin, R.E., Savic, G., Chang, J., Carson, M.B., Primeaux, C., Tian, R., Kenton, S., Jia, H., et al. (2002). Genome sequence of Streptococcus mutans UA159, a cariogenic dental pathogen. Proc Natl Acad Sci U S A 99, 14434–14439. https://doi.org/10.1073/pnas.172501299.

Alvarez-Martinez, C.E., and Christie, P.J. (2009). Biological diversity of prokaryotic Type IV secretion systems. Microbiol Mol Bio Rev 73, 775–808. https://doi.org/10.1128/MMBR.00023-09.

Auchtung, J.M., Lee, C.A., Monson, R.E., Lehman, A.P., and Grossman, A.D. (2005). Regulation of a Bacillus subtilis mobile genetic element by intercellular signaling and the global DNA damage response. Proc Natl Acad Sci U S A 102, 12554–12559. https://doi.org/10.1073/pnas.0505835102.

Auchtung, J.M., Lee, C.A., Garrison, K.L., and Grossman, A.D. (2007). Identification and characterization of the immunity repressor (ImmR) that controls the mobile genetic element ICEBs1 of Bacillus subtilis. Mol Microbiol 64, 1515–1528. https://doi.org/10.1111/j.1365-2958.2007.05748.x.

Auchtung, J.M., Aleksanyan, N., Bulku, A., and Berkmen, M.B. (2016). Biology of ICEBs1, an integrative and conjugative element in Bacillus subtilis. Plasmid 86, 14–25. https://doi.org/10.1016/j.plasmid.2016.07.001.

Babic, A., Berkmen, M.B., Lee, C.A., and Grossman, A.D. (2011). Efficient gene transfer in bacterial cell chains. mBio 2, e00027–11. https://doi.org/10.1128/mBio.00027-11.

Beaber, J.W., Hochhut, B., and Waldor, M.K. (2004). SOS response promotes horizontal dissemination of antibiotic resistance genes. Nature 427, 72–74. https://doi.org/10.1038/nature02241.

Bean, E.L., McLellan, L.K., and Grossman, A.D. (2022a). Activation of the integrative and conjugative element Tn916 causes growth arrest and death of host bacteria. bioRvix 2022.04.01.486793. https://doi.org/10.1101/2022.04.01.486793.

Bean, E.L., Herman, C., Anderson, M.E., and Grossman, A.D. (2022b). Biology and engineering of integrative and conjugative elements: Construction and analyses of hybrid ICEs reveal element functions that affect species-specific efficiencies. PLoS Genet 18, e1009998. https://doi.org/10.1371/journal.pgen.1009998.

Benson, A.K., and Haldenwang, W.G. (1993). Regulation of sigma B levels and activity in Bacillus subtilis. J Bacteriol 175, 2347–2356.

Berkmen, M.B., Lee, C.A., Loveday, E.-K., and Grossman, A.D. (2010). Polar positioning of a conjugation protein from the integrative and conjugative element ICEBs1 of Bacillus subtilis. J Bacteriol 192, 38–45. https://doi.org/10.1128/JB.00860-09.

Bhatty, M., Laverde Gomez, J.A., and Christie, P.J. (2013). The expanding bacterial type IV secretion lexicon. Res Microbiol 164, 620–639. https://doi.org/10.1016/j.resmic.2013.03.012.

Bi, D., Xu, Z., Harrison, E.M., Tai, C., Wei, Y., He, X., Jia, S., Deng, Z., Rajakumar, K., and Ou, H.-Y. (2012). ICEberg: a web-based resource for integrative and conjugative elements found in bacteria. Nucleic Acids Res 40, D621–D626. https://doi.org/10.1093/nar/gkr846.

Bose, B., and Grossman, A.D. (2011). Regulation of horizontal gene transfer in Bacillus subtilis by activation of a conserved site-specific protease. J Bacteriol 193, 22–29. https://doi.org/10.1128/JB.01143-10.

Bose, B., Auchtung, J.M., Lee, C.A., and Grossman, A.D. (2008). A conserved anti-repressor controls horizontal gene transfer by proteolysis. Mol Microbiol 70, 570–582. https://doi.org/10.1111/j.1365-2958.2008.06414.x.

Botelho, J., and Schulenburg, H. (2021). The role of integrative and conjugative elements in antibiotic resistance evolution. Trends Microbiol 29, 8–18. https://doi.org/10.1016/j.tim.2020.05.011.

Brophy, J.A.N., Triassi, A.J., Adams, B.L., Renberg, R.L., Stratis-Cullum, D.N., Grossman, A.D., and Voigt, C.A. (2018). Engineered integrative and conjugative elements for efficient and inducible DNA transfer to undomesticated bacteria. Nat Microbiol 3, 1043–1053. https://doi.org/10.1038/s41564-018-0216-5.

Burrus, V., and Waldor, M.K. (2004). Shaping bacterial genomes with integrative and conjugative elements. Rese Microbiol 155, 376–386. https://doi.org/10.1016/j.resmic.2004.01.012.

Burrus, V., Pavlovic, G., Decaris, B., and Guédon, G. (2002). Conjugative transposons: the tip of the iceberg. Mol Microbiol 46, 601–610. https://doi.org/10.1046/j.1365-2958.2002.03191.x.

Ciric, L., Jasni, A., Vries, L.E. de Agersø, Y., Mullany, P., and Roberts, A.P. (2013). The Tn916/Tn1545 ramily of conjugative transposons (Landes Bioscience).

Cury, J., Touchon, M., and Rocha, E.P.C. (2017). Integrative and conjugative elements and their hosts: composition, distribution and organization. Nucleic Acids Res 45, 8943–8956. https://doi.org/10.1093/nar/gkx607.

Decker, E.-M. (2001). The ability of direct fluorescence-based, two-colour assays to detect different physiological states of oral streptococci. Lett Appl Microbiol 33, 188–192. https://doi.org/10.1046/j.1472-765x.2001.00971.x.

Decker, E.-M., Klein, C., Schwindt, D., and von Ohle, C. (2014). Metabolic activity of Streptococcus mutans biofilms and gene expression during exposure to xylitol and sucrose. Int J Oral Sci 6, 195–204. https://doi.org/10.1038/ijos.2014.38.

Delavat, F., Mitri, S., Pelet, S., and van der Meer, J.R. (2016). Highly variable individual donor cell fates characterize robust horizontal gene transfer of an integrative and conjugative element. Proc Natl Acad Sci USA 113, E3375–E3383. https://doi.org/10.1073/pnas.1604479113.

Dimopoulou, I.D., Russell, J.E., Mohd-Zain, Z., Herbert, R., and Crook, D.W. (2002). Site-specific recombination with the chromosomal tRNA(Leu) gene by the large conjugative Haemophilus resistance plasmid. Antimicrob Agents Chemother 46, 1602–1603. https://doi.org/10.1128/AAC.46.5.1602-1603.2002.

Franke, A.E., and Clewell, D.B. (1981a). Evidence for a chromosome-borne resistance transposon (Tn916) in Streptococcus faecalis that is capable of “conjugal” transfer in the absence of a conjugative plasmid. J Bacteriol 145, 494–502. https://doi.org/10.1128/jb.145.1.494-502.1981.

Franke, A.E., and Clewell, D.B. (1981b). Evidence for conjugal transfer of a Streptococcus faecalis transposon (Tn916) from a chromosomal site in the absence of plasmid DNA. Cold Spring Harb Symp Quant Biol 45 Pt 1, 77–80. https://doi.org/10.1101/sqb.1981.045.01.014.

Fronzes, R., Christie, P.J., and Waksman, G. (2009). The structural biology of type IV secretion systems. Nat Rev Microbiol 7, 10.1038/nrmicro2218. https://doi.org/10.1038/nrmicro2218.

Fujimura-Kamada, K., Nouvet, F.J., and Michaelis, S. (1997). A novel membrane-associated metalloprotease, Ste24p, is required for the first step of NH2-terminal processing of the yeast a-factor precursor. J Cell Biol 136, 271–285. https://doi.org/10.1083/jcb.136.2.271.

Gao, Y., Mathee, K., Narasimhan, G., and Wang, X. (1999). Motif detection in protein sequences. In 6th International Symposium on String Processing and Information Retrieval. 5th International Workshop on Groupware. PR00268 (IEEE), pp. 63-72.

Gibson, D. (2009). One-step enzymatic assembly of DNA molecules up to several hundred kilobases in size. Protoc Exch https://doi.org/10.1038/nprot.2009.77.

Gilchrist, C.L.M., Booth, T.J., van Wersch, B., van Grieken, L., Medema, M.H., and Chooi, Y.-H. (2021). cblaster: a remote search tool for rapid identification and visualization of homologous gene clusters. Bioinform Adv 1, vbab016. https://doi.org/10.1093/bioadv/vbab016.

Gottesman, M.E., and Weisberg, R.A. (2004). Little lambda, who made thee? Microbiol Mol Biol Rev 68, 796–813. https://doi.org/10.1128/MMBR.68.4.796-813.2004.

Grindley, N.D.F., Whiteson, K.L., and Rice, P.A. (2006). Mechanisms of site-specific recombination. Ann Rev Biochem 75, 567–605. https://doi.org/10.1146/annurev.biochem.73.011303.073908.

Guérout-Fleury, A.M., Frandsen, N., and Stragier, P. (1996). Plasmids for ectopic integration in Bacillus subtilis. Gene 180, 57–61. https://doi.org/10.1016/s0378-1119(96)00404-0.

Guglielmini, J., Quintais, L., Garcillán-Barcia, M.P., de la Cruz, F., and Rocha, E.P.C. (2011). The repertoire of ICE in prokaryotes underscores the unity, diversity, and ubiquity of conjugation. PLoS Genet 7, e1002222. https://doi.org/10.1371/journal.pgen.1002222.

Harwood, C.R., and Cutting, S.M. (1990). Molecular biological methods for Bacillus (Wiley).

Hirano, N., Muroi, T., Takahashi, H., and Haruki, M. (2011). Site-specific recombinases as tools for heterologous gene integration. Appl Microbiol Biotechnol 92, 227–239. https://doi.org/10.1007/s00253-011-3519-5.

Iyer, L.M., Makarova, K.S., Koonin, E.V., and Aravind, L. (2004). Comparative genomics of the FtsK–HerA superfamily of pumping ATPases: implications for the origins of chromosome segregation, cell division and viral capsid packaging. Nucleic Acids Res 32, 5260–5279. https://doi.org/10.1093/nar/gkh828.

Johnson, C.M., and Grossman, A.D. (2014). Identification of host genes that affect acquisition of an integrative and conjugative element in Bacillus subtilis. Mol Microbiol 93, 1284–1301. https://doi.org/10.1111/mmi.12736.

Johnson, C.M., and Grossman, A.D. (2015). Integrative and conjugative elements (ICEs): what they do and how they work. Annu Rev Genet 49, 577–601. https://doi.org/10.1146/annurev-genet-112414-055018.

Jones, J.M., Grinberg, I., Eldar, A., and Grossman, A.D. (2021). A mobile genetic element increases bacterial host fitness by manipulating development. eLife 10, e65924. https://doi.org/10.7554/eLife.65924.

King, S., Quick, A., King, K., Walker, A.R., and Shields, R.C. (2022). Activation of TnSmu1, an integrative and conjugative element, is lethal to Streptococcus mutans. bioRvix 2022.05.11.491493. https://doi.org/10.1101/2022.05.11.491493.

Koyanagi, S., and Lévesque, C.M. (2013). Characterization of a Streptococcus mutans intergenic region containing a small toxic peptide and its cis-encoded antisense small RNA antitoxin. PLoS ONE 8, e54291. https://doi.org/10.1371/journal.pone.0054291.

Lee, C.A., and Grossman, A.D. (2007). Identification of the origin of transfer (oriT) and DNA relaxase required for conjugation of the integrative and conjugative element ICEBs1 of Bacillus subtilis. J Bacteriol 189, 7254–7261. https://doi.org/10.1128/JB.00932-07.

Lee, C.A., Auchtung, J.M., Monson, R.E., and Grossman, A.D. (2007). Identification and characterization of int (integrase), xis (excisionase) and chromosomal attachment sites of the integrative and conjugative element ICEBs1 of Bacillus subtilis. Mol Microbiol 66, 1356–1369. https://doi.org/10.1111/j.1365-2958.2007.06000.x.

Lee, C.A., Babic, A., and Grossman, A.D. (2010). Autonomous plasmid-like replication of a conjugative transposon. Mol Microbiol 75, 268–279. https://doi.org/10.1111/j.1365-2958.2009.06985.x.

Lee, C.A., Thomas, J., and Grossman, A.D. (2012). The Bacillus subtilis conjugative transposon ICEBs1 mobilizes plasmids lacking dedicated mobilization functions. J Bacteriol 194, 3165– 3172. https://doi.org/10.1128/JB.00301-12.

Lemon, K.P., Kurtser, I., and Grossman, A.D. (2001). Effects of replication termination mutants on chromosome partitioning in Bacillus subtilis. Proc Natl Acad Sci U S A 98, 212–217.

Leonetti, C.T., Hamada, M.A., Laurer, S.J., Broulidakis, M.P., Swerdlow, K.J., Lee, C.A., Grossman, A.D., and Berkmen, M.B. (2015). Critical components of the conjugation machinery of the integrative and conjugative element ICEBs1 of Bacillus subtilis. J Bacteriol 197, 2558– 2567. https://doi.org/10.1128/JB.00142-15.

Li, Y.-H., Lau, P.C.Y., Lee, J.H., Ellen, R.P., and Cvitkovitch, D.G. (2001). Natural genetic transformation of Streptococcus mutans growing in biofilms. J Bacteriol 183, 897–908. https://doi.org/10.1128/JB.183.3.897-908.2001.

Lucchini, S., Desiere, F., and Brüssow, H. (1999). Similarly organized lysogeny modules in temperate Siphoviridae from low GC content gram-positive bacteria. Virology 263, 427–435. https://doi.org/10.1006/viro.1999.9959.

Lunde, T.M., Hjerde, E., and Al-Haroni, M. (2021). Prevalence, diversity and transferability of the Tn 916 -Tn 1545 family ICE in oral streptococci. J Oral Microbiol 13, 1896874. https://doi.org/10.1080/20002297.2021.1896874.

Mashburn-Warren, L., Morrison, D.A., and Federle, M.J. (2010). A novel double-tryptophan peptide pheromone is conserved in mutans and pyogenic Streptococci and Controls Competence in Streptococcus mutans via an Rgg regulator. Mol Microbiol 78, 589–606. https://doi.org/10.1111/j.1365-2958.2010.07361.x.

Miyazaki, R., and van der Meer, J.R. (2013). A new large-DNA-fragment delivery system based on integrase activity from an integrative and conjugative element. Appl Environ Microbiol 79, 4440–4447. https://doi.org/10.1128/AEM.00711-13.

Narasimhan, G., Bu, C., Gao, Y., Wang, X., Xu, N., and Mathee, K. (2002). Mining protein sequences for motifs. J Comput Biol 9, 707–720. https://doi.org/10.1089/106652702761034145.

Needleman, S.B., and Wunsch, C.D. (1970). A general method applicable to the search for similarities in the amino acid sequence of two proteins. J Mol Biol 48, 443–453. https://doi.org/10.1016/0022-2836(70)90057-4.

Nomura, R., Matayoshi, S., Otsugu, M., Kitamura, T., Teramoto, N., and Nakano, K. Contribution of severe dental caries induced by Streptococcus mutans to the pathogenicity of infective endocarditis. Infect Immun 88, e00897–19. https://doi.org/10.1128/IAI.00897-19.

Olsen, I., Tribble, G.D., Fiehn, N.-E., and Wang, B.-Y. (2013). Bacterial sex in dental plaque. J Oral Microbiol 5, 10.3402/jom.v5i0.20736. https://doi.org/10.3402/jom.v5i0.20736.

Oppenheim, A.B., Kobiler, O., Stavans, J., Court, D.L., and Adhya, S. (2005). Switches in bacteriophage lambda development. Annu Rev of Genet 39, 409–429. https://doi.org/10.1146/annurev.genet.39.073003.113656.

Pembroke, J.T., and Stevens, E. (1984). The effect of plasmid R391 and other IncJ plasmids on the survival of Escherichia coli after UV irradiation. J Gen Microbiol 130, 1839–1844. https://doi.org/10.1099/00221287-130-7-1839.

Peters, J.M., Koo, B.-M., Patino, R., Heussler, G.E., Hearne, C.C., Qu, J., Inclan, Y.F., Hawkins, J.S., Lu, C.H.S., Silvis, M.R., et al. (2019). Enabling genetic analysis of diverse bacteria with Mobile-CRISPRi. Nat Microbiol 4, 244–250. https://doi.org/10.1038/s41564-018-0327-z.

Petersen, F.C., and Scheie, A.A. (2010). Natural transformation of oral streptococci. In Oral Biology: Molecular Techniques and Applications, G.J. Seymour, M.P. Cullinan, and N.C.K. Heng, eds. (Totowa, NJ: Humana Press), pp. 167–180.

Petit, M.-A., Dervyn, E., Rose, M., Entian, K.-D., McGovern, S., Ehrlich, S.D., and Bruand, C. (1998). PcrA is an essential DNA helicase of Bacillus subtilis fulfilling functions both in repair and rolling-circle replication. Mol Microbiol 29, 261–273. https://doi.org/10.1046/j.1365-2958.1998.00927.x.

Pfaffl, M.W. (2001). A new mathematical model for relative quantification in real-time RT-PCR. Nucleic Acids Res 29, e45. https://doi.org/10.1093/nar/29.9.e45.

Reck, M., and Wagner-Döbler, I. (2016). Carolacton treatment causes delocalization of the cell division proteins PknB and DivIVa in Streptococcus mutans in vivo. Front Microbiol 7.

Reinhard, F., Miyazaki, R., Pradervand, N., and van der Meer, J.R. (2013). Cell differentiation to “mating bodies” induced by an integrating and conjugative element in free-living bacteria. Curr Biol 23, 255–259. https://doi.org/10.1016/j.cub.2012.12.025.

Rice, P., Longden, I., and Bleasby, A. (2000). EMBOSS: the European Molecular Biology Open Software Suite. Trends Genet 16, 276–277. https://doi.org/10.1016/s0168-9525(00)02024-2.

Roberts, A.P., and Mullany, P. (2009). A modular master on the move: the Tn916 family of mobile genetic elements. Trends Microbiol 17, 251–258. https://doi.org/10.1016/j.tim.2009.03.002.

Roberts, A.P., and Mullany, P. (2011). Tn916-like genetic elements: a diverse group of modular mobile elements conferring antibiotic resistance. FEMS Microbiol Rev 35, 856–871. https://doi.org/10.1111/j.1574-6976.2011.00283.x.

Roberts, A.P., Pratten, J., Wilson, M., and Mullany, P. (1999). Transfer of a conjugative transposon, Tn5397 in a model oral biofilm. FEMS Microbiol Lett 177, 63–66. https://doi.org/10.1111/j.1574-6968.1999.tb13714.x.

Roberts, A.P., Cheah, G., Ready, D., Pratten, J., Wilson, M., and Mullany, P. (2001). Transfer of Tn916-like elements in microcosm dental plaques. Antimicrob Agents Chemother 45, 2943– 2946. https://doi.org/10.1128/AAC.45.10.2943-2946.2001.

Rocco, J.M., and Churchward, G. (2006). The integrase of the conjugative transposon Tn916 directs strand- and sequence-specific cleavage of the origin of conjugal transfer, oriT, by the endonuclease Orf20. J Bacteriol 188, 2207–2213. https://doi.org/10.1128/JB.188.6.2207-2213.2006.

San Millan, A., and MacLean, R.C. (2017). Fitness costs of plasmids: a limit to plasmid transmission. Microbiol Spectr 5, 5.5.02. https://doi.org/10.1128/microbiolspec.MTBP-0016-2017.

Santoro, F., Vianna, M.E., and Roberts, A.P. (2014). Variation on a theme; an overview of the Tn916/Tn1545 family of mobile genetic elements in the oral and nasopharyngeal streptococci. Front Microbiol 5, 535. https://doi.org/10.3389/fmicb.2014.00535.

Serfiotis-Mitsa, D., Roberts, G.A., Cooper, L.P., White, J.H., Nutley, M., Cooper, A., Blakely, G.W., and Dryden, D.T.F. (2008). The Orf18 gene product from conjugative transposon Tn916 is an ArdA antirestriction protein that inhibits type I DNA restriction–modification systems. J Mol Biol 383, 970–981. https://doi.org/10.1016/j.jmb.2008.06.005.

Shanker, E., and Federle, M.J. (2017). Quorum sensing regulation of competence and bacteriocins in Streptococcus pneumoniae and mutans. Genes 8, 15. https://doi.org/10.3390/genes8010015.

Shields, R.C., Zeng, L., Culp, D.J., and Burne, R.A. (2018). Genomewide identification of essential genes and fitness determinants of Streptococcus mutans UA159. mSphere 3, e00031–18. https://doi.org/10.1128/mSphere.00031-18.

Son, M., Ahn, S.-J., Guo, Q., Burne, R.A., and Hagen, S.J. (2012). Microfluidic study of competence regulation in Streptococcus mutans: environmental inputs modulate bimodal and unimodal expression of comX. Mol Microbiol 86, 258–272. https://doi.org/10.1111/j.1365-2958.2012.08187.x.

Thomas, J., Lee, C.A., and Grossman, A.D. (2013). A conserved helicase processivity factor is needed for conjugation and replication of an integrative and conjugative element. PLoS Genet 9. https://doi.org/10.1371/journal.pgen.1003198.

Wecke, J., Madela, K., and Fischer, W. (1997). The absence of D-alanine from lipoteichoic acid and wall teichoic acid alters surface charge, enhances autolysis and increases susceptibility to methicillin in Bacillus subtilis. Microbiol 143, 2953–2960. https://doi.org/10.1099/00221287-143-9-2953.

Wozniak, R.A.F., and Waldor, M.K. (2010). Integrative and conjugative elements: mosaic mobile genetic elements enabling dynamic lateral gene flow. Nat Rev Microbiol 8, 552–563. https://doi.org/10.1038/nrmicro2382.

Wright, L.D., and Grossman, A.D. (2016). Autonomous replication of the conjugative transposon Tn916. J Bacteriol. 198, 3355–3366. https://doi.org/10.1128/JB.00639-16.

Xiang, Y., Morais, M.C., Cohen, D.N., Bowman, V.D., Anderson, D.L., and Rossmann, M.G. (2008). Crystal and cryoEM structural studies of a cell wall degrading enzyme in the bacteriophage phi29 tail. Proc Natl Acad Sci U S A 105, 9552–9557. https://doi.org/10.1073/pnas.0803787105.

Xie, Z., Okinaga, T., Qi, F., Zhang, Z., and Merritt, J. (2011). Cloning-independent and counterselectable markerless mutagenesis system in Streptococcus mutans. Appl Environ Microbiol 77, 8025–8033. https://doi.org/10.1128/AEM.06362-11.

Zhang, K., Ou, M., Wang, W., and Ling, J. (2009). Effects of quorum sensing on cell viability in Streptococcus mutans biofilm formation. Biochem Biophys Res Communs 379, 933–938. https://doi.org/10.1016/j.bbrc.2008.12.175.

